# The genomic variation landscape of globally-circulating clades of SARS-CoV-2 defines a genetic barcoding scheme

**DOI:** 10.1101/2020.04.21.054221

**Authors:** Qingtian Guan, Mukhtar Sadykov, Raushan Nugmanova, Michael J. Carr, Stefan T. Arold, Arnab Pain

## Abstract

We describe fifteen major mutation events from 2,058 high-quality SARS-CoV-2 genomes deposited up to March 31^st^, 2020. These events define five major clades (G, I, S, D and V) of globally-circulating viral populations, representing 85.7% of all sequenced cases, which we can identify using a 10 nucleotide genetic classifier or barcode. We applied this barcode to 4,000 additional genomes deposited between March 31^st^ and April 15^th^ and classified successfully 95.6% of the clades demonstrating the utility of this approach. An analysis of amino acid variation in SARS-CoV-2 ORFs provided evidence of substitution events in the viral proteins involved in both host-entry and genome replication. The systematic monitoring of dynamic changes in the SARS-CoV-2 genomes of circulating virus populations over time can guide therapeutic and prophylactic strategies to manage and contain the virus and, also, with available efficacious antivirals and vaccines, aid in the monitoring of circulating genetic diversity as we proceed towards elimination of the agent. The barcode will add the necessary genetic resolution to facilitate tracking and monitoring of infection clusters to distinguish imported and indigenous cases and thereby aid public health measures seeking to interrupt transmission chains without the requirement for real-time complete genomes sequencing.

## INTRODUCTION

SARS-CoV-2 has now reached 185 countries across all continents, except Antarctica. With over 2.5 million cases currently confirmed globally^1^, a pandemic was declared by the World Health Organisation (WHO) on March 11^th^, 2020. SARS-CoV-2 belongs to the Coronaviridae family, genus Betacoronavirus, which are enveloped positive-sense, single-stranded RNA viruses, of zoonotic origin. Among human RNA viruses, coronaviruses have the largest known genome (∼30 kb), which consists of the structural proteins (spike, envelope, membrane and nucleocapsid), nonstructural proteins (nsp1-16), and accessory proteins (ORF3a, ORF6, ORF7a and b, ORF8, ORF10). Structural proteins are required for host cell entry, viral assembly and exit^2,3^. Nonstructural proteins are involved in genome replication-transcription and formation of vesicles^4^, whereas accessory proteins interfere with host innate defense mechanisms^5,6,7^. During replication within the host, the virus acquires genome mutations, which can be passed on to virus progeny in subsequently infected individuals. Systematically tracking mutations in SARS-CoV-2 genomes is therefore important as it allows monitoring of the molecular epidemiology of circulating viral sequences nationally and internationally. Here, we have investigated the genomic variation landscape of a large set of globally-derived SARS-CoV-2 genomes and defined major mutation events. This analysis allowed us to produce a first-generation genetic classifier, or ‘barcode’, defining the major clades of the virus circulating up to April 15^th^, 2020. Notably, this barcode allowed reliable tracking of the spatial distribution and prevalence of these viral clades over time. While most of the nonsynonymous mutation events appear neutral with respect to protein function and stability, we also found evidence of mutations in the spike protein that may modulate the interaction between SARS-CoV-2 and the host.

## RESULTS

### Five clades of SARS-CoV-2 are characterised by 15 major mutational events across the globe

The SARS-CoV-2 genome is genetically most closely related (96%) to a bat SARS-related coronavirus (SARSr-CoV) RaTG13 and also to SARSr-CoVs from pangolins^8^ (Supplementary Figure 1). We analysed single nucleotide polymorphisms (SNPs) of 2,058 high-quality complete genomes downloaded from GISAID^9^ (March 31^st^). We studied their chronological occurrences during the spread of the SARS-CoV-2 across human populations, which allowed us to define several major and minor clades of the virus that share unique SNPs (Supplementary Table 1, Supplementary Figures 2, 3). We observed 1,221 SNPs in the current dataset with 753 missense, 452 silent, 12 nonsense and 4 intergenic substitutions. We defined five major clades (Figure 1) compared to the prototype (MN908947.3), covering 85.7% of the global set of SARS-CoV-2 high-quality genomes publicly available up to March 31^st^. The clades were named by the amino acid mutation: S (*Orf8*, L84S), V (*Orf3a*, G251V), I (*Orf1ab*, V378I), D (*Orf1ab*, G392D) and G (*S*, D614G).

**Table 1.**
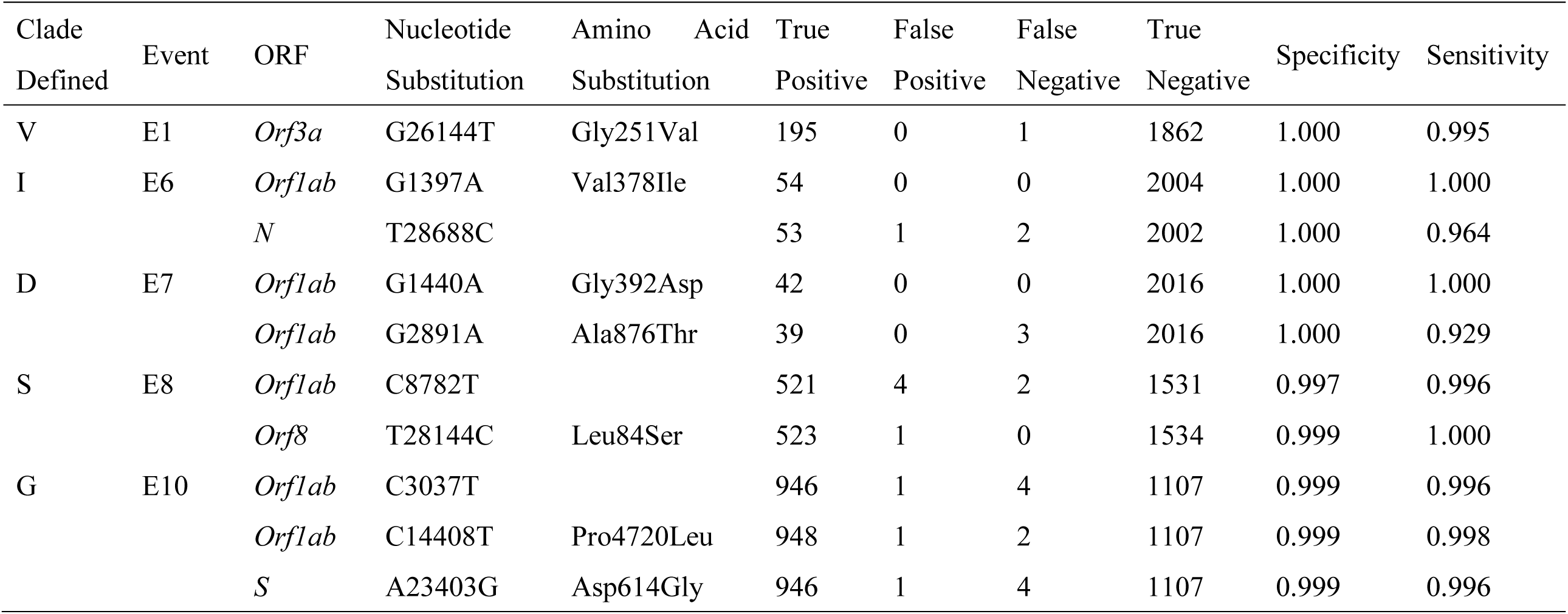
Sensitivity and specificity of the SARS-CoV-2 clade-defining SNP based on 2,058 genomes covering the first 14 weeks of the COVID-19 outbreak.

**Figure 1.**
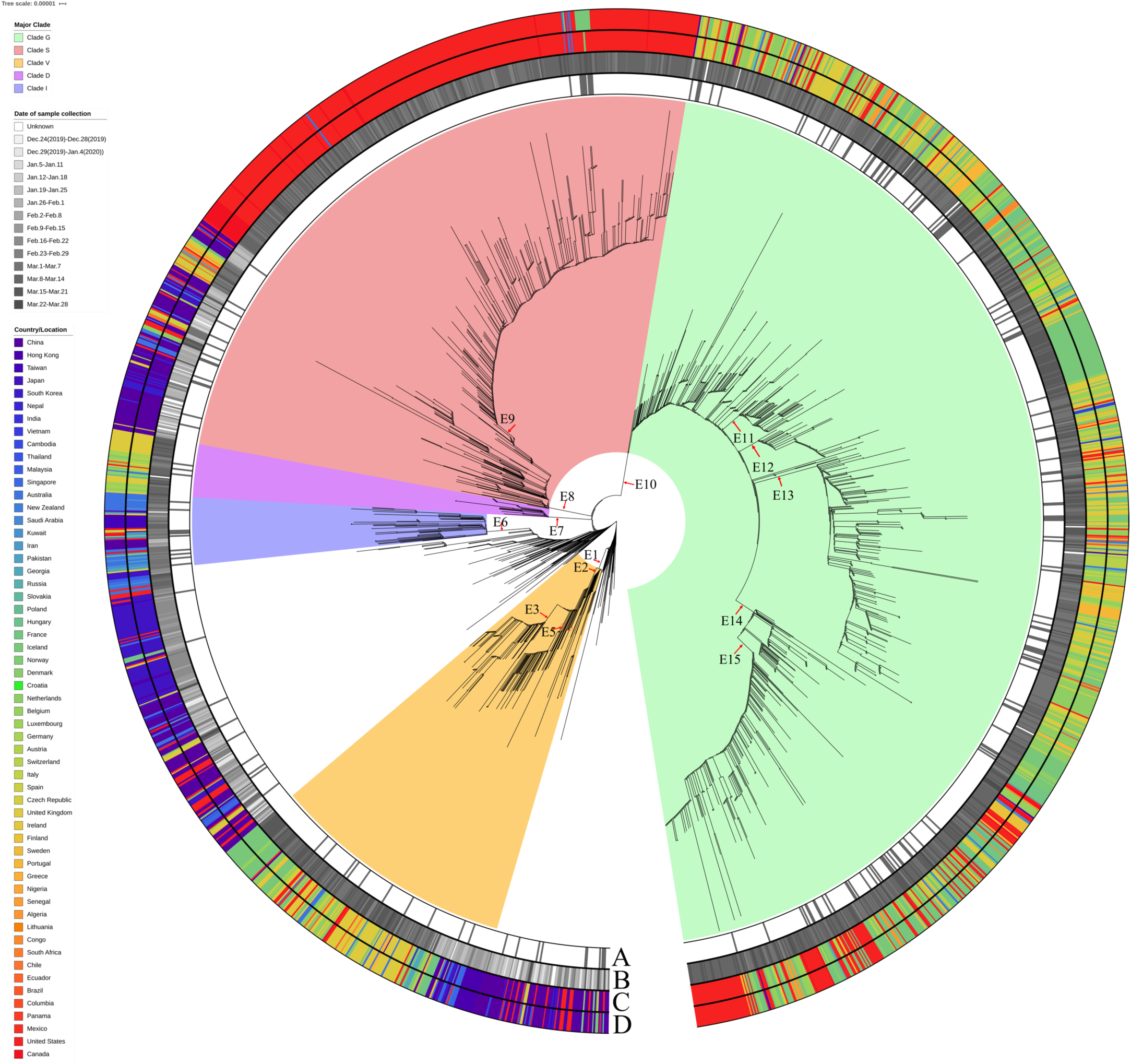
Major substitutions events in SARS-CoV-2. A global SNP-based radial phylogeny of SARS-CoV-2 genomes defining five major clades (S, G, V, D and I) and several subclades based on nucleotide substitution events. E1-E15 represents the 15 major evolutionary events. Branches with >70% bootstrap values are shown and all the main clade-defining branches have bootstrap values >90%. E4, which corresponds to a G11083A mutation in ORF1a was found in two independent branches of the tree and for illustration purposes it is shown in Supplementary Figure 2. The labeling for the concentric outer circles I as follows: A, the imported cases which the country of exposure differs from the country of isolation; B, the collection date of each case with one week resolution; C, collection locations of the cases; D exposure locations of the cases.

These 5 clades are characterised by 15 major nucleotide substitution events in the SARS-CoV-2 genome (Figure 1, Figure 2 and Supplementary Figure 3) representing 1,763 genomes (85.7%) from 45 countries. To obtain a global picture of the regional distribution of the clades over time during a 14 week period, we plotted the relative proportions of the major and minor clades and their cumulative trend (Figure 3). We observed major differences in the apparent spread for individual clades: Clade G represents 46.2% of all the sequenced viral sequences, followed by S (25.4%), V (9.4%), I (2.6%) and D (2.1%). The remaining 14.3% were not assigned to a major clade. Clade G is widely distributed in Africa, Europe, West Asia and South America; whereas Clade S represents 63% of North American sampled genomes, and nearly a quarter of those from Oceania. Clade I represents around one-third of genomes derived from South and West Asia, and Oceania. Southeast Asia and South Asia have the greatest number of unassigned genomes (56.9%). For these genomes that cannot be assigned to a major clade, we identified nine minor clades which were named for the amino acid mutation or nucleotide substitution (the latter shown in bold): H (*Orf1ab*, Q676H); H2 (*M*, D209H); L2 (*N*, S194L, we name it L2 to avoid confusion with the previously defined clade^8^); S2 (*N*, P344S); G11410A (*Orf1ab*, **G11410A**); Y (*S*, H49Y); C17373A (*Orf1ab*, **C17373A**); I2 (*Orf1ab*, T6136I) and K (*Orf1ab*, T2016K). The minor clades represent any monoclades with n ≥ 5 which covers 3.2% of the total 2,058 cases. The global and regional cumulative trends were plotted over time, with the majority of the trends revealing the increasing dominance of one or two clades in each geographic region. For example, the Asian and Oceanian genomes are largely clade I whereas European genomes are predominantly clade G with clade S predominating in North America cases. This is likely attributable to founder effects during the early phases of the seeding of the local epidemics from imported cases and subsequent dissemination in the regions.

**Figure 2.**
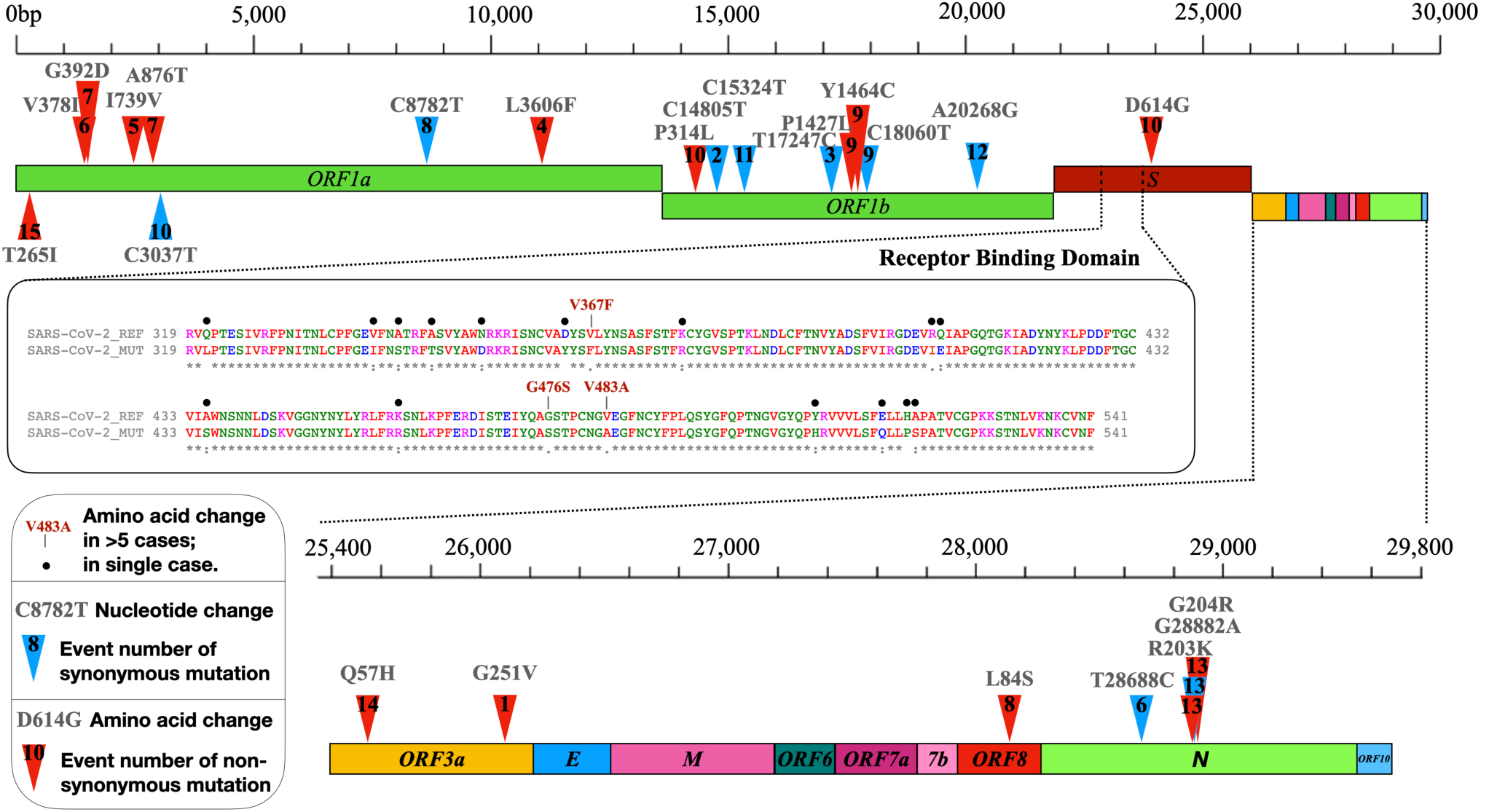
Major mutations and associated variation in globally circulating SARS-CoV-2 genomes (n=2,058). Genomic localisation of major mutation events as defined within our study. SARS-CoV-2 mutations in the receptor-bing domain (RBD) sequence contain all amino acid substitutions from 2,058 genomes available up until March 31^st^ 2020.

**Figure 3.**
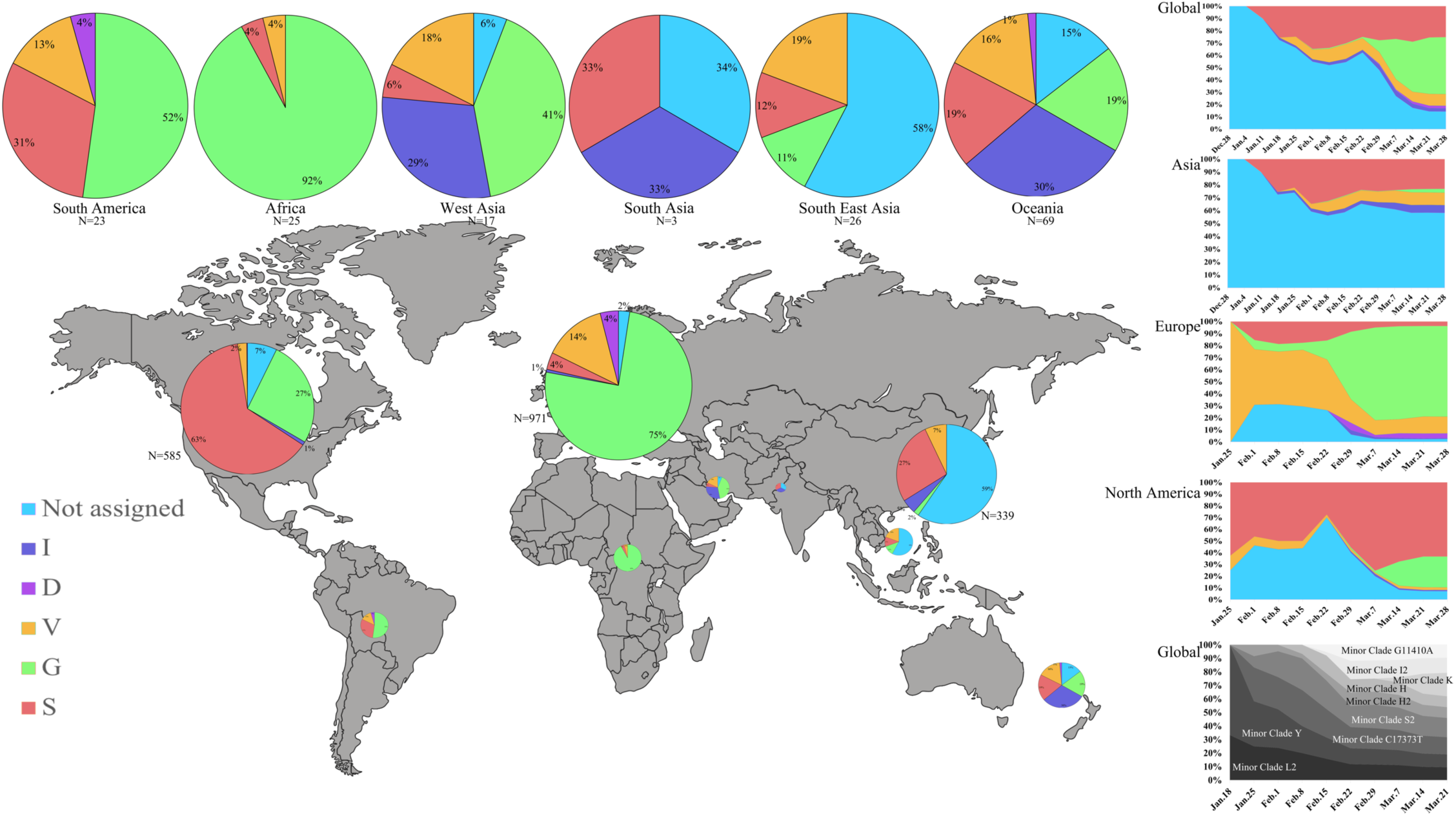
Global distribution of various major and minor clades of SARS-CoV2 genomes and their relative prevalence over a 14 week time period from December 24^th^ 2019 to March 28^th^ 2020 from the outbreak and early stages of the pandemic. The size of each pie chart is proportional to the numbers within each respective clade (Europe, max=971; South Asia, min=3). The cumulative trend of the clades is shown on the right and the span of time indicates the first and last observed case in each particular clade, until March 28^th^ 2020.

### The genetic barcoding method assigns new SARS-CoV-2 cases to clades with high sensitivity

We defined a 10 nucleotide genetic barcode of SARS-CoV-2 that identified with high sensitivity the five major clades of the circulating viral genomes available on March 31^st^, based on the GISAID data (Figure 4A, Table 1). Given that SARS-CoV-2 evolves at an average rate/genome of nearly 8 × 10^−4^ nucleotide substitutions/site/year^10^, and is subject to back-mutations, we further tested the sensitivity and specificity of these clade-defining SNPs, which are all above 90% (Table 1). We then applied the 10 nucleotide barcode to the 4,000 SARS-CoV-2 genomes that became available in GISAID between March 31^st^ and April 15^th^, 2020 in the early stages of the pandemic. Among the 4,000 globally-circulating genomes from 66 countries, we could assign ∼96% to one of the 5 major clades (Figure 4B). The remaining unassigned 4% of genomes were typified by either errors in their sequences (such as ‘N’s in the genome assemblies), or single cases those could not be reliably assigned to any of the major clades. The increase in the correct clade assignment for the new genomes compared to the initial validation of the methodology on 2,058 genomes (from which the barcode was established) was indicative of the rising dominance of a few major clades, and suggested that the 10 nucleotide barcode will retain its predictive power as new genomes are obtained. While this barcode represents a snapshot of the early phases of the genetic diversity of the virus during the first 16 weeks of its global spread and is expected to change over time, a barcoding strategy to monitor the progress of virus elimination after vaccines become widely available will be strategically useful to monitor decreases in viral genetic diversity. In addition, our barcode could serve as a reference for setting the baseline for global genomic diversity analysis at the beginning of the pandemic.

**Figure 4.**
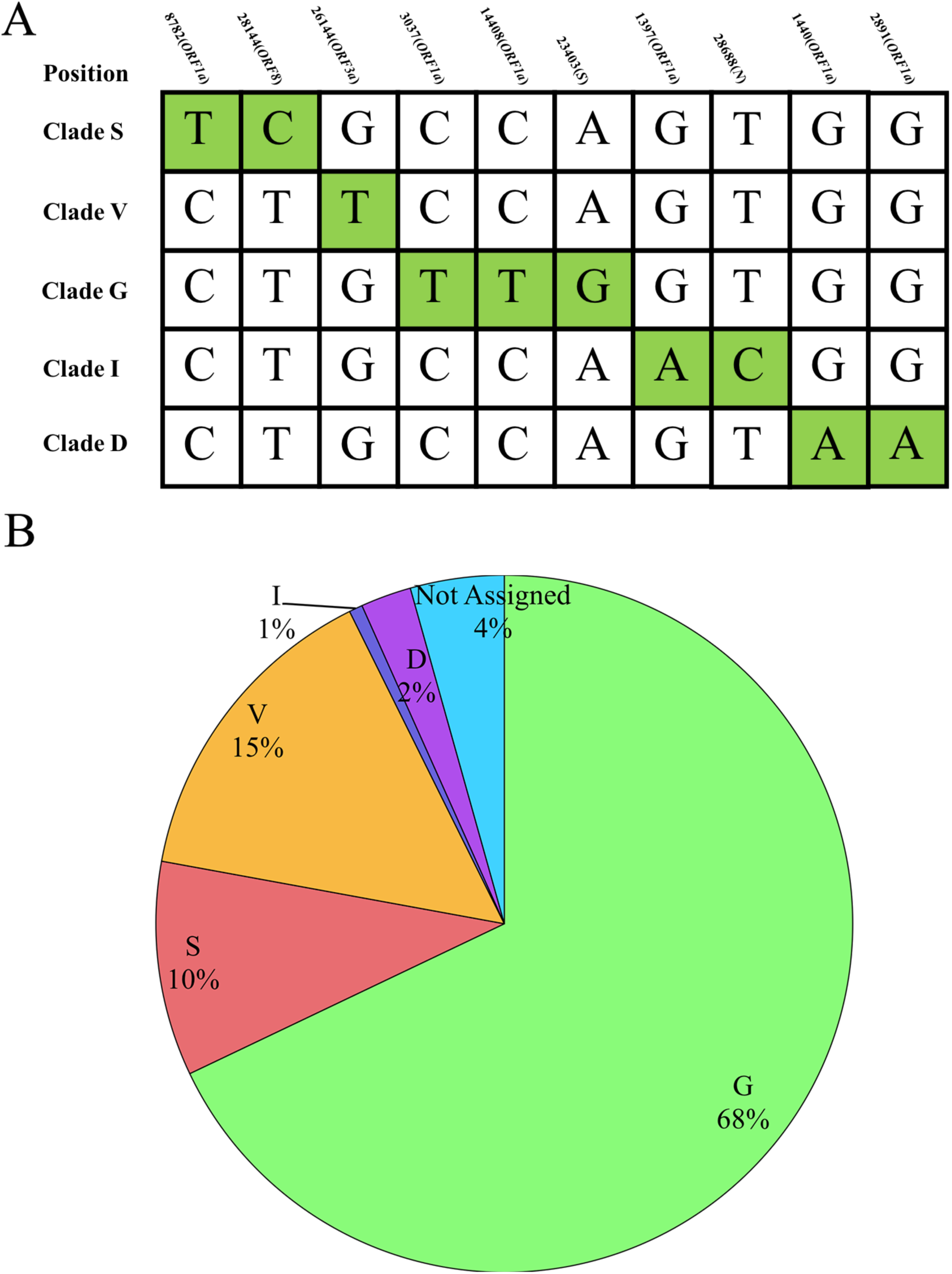
Major mutations and associated variation in globally circulating SARS-CoV-2 genomes (n=6,058) available until April 15^th^ 2020. (A) A 10 nucleotide SNP genetic barcode that defines the 5 clades of SARS-CoV-2. (B) By applying this barcode we are able to classify 95.6% of the cases published between March 31^st^ and April 15^th^ 2020 during the early phase of the pandemic. Pie charts showing the percentage of each major clade by applying a 10 nucleotide genetic classifier to 4,000 new genomes downloaded from GISAID.

### Mutations are not equally distributed across the SARS-CoV-2 genome

Based on our analysis of the 2,058 available genomes from GISAID (March 31^th^, 2020) we observed that the genes *S, N* and *Orf3a*, accumulated markedly more mutations than expected solely by random drift (Supplementary Figure 4) (Real/Expected ratio: *S*: 1.21; *N*: 1.99; *ORF3a*: 1.82). This mutation rate may indicate adaptation to the human host following recent spill over from an, as yet unknown, animal reservoir. Conversely, several nonstructural proteins showed a lower-than-expected mutation rate (Real/Expected ratio: nsp1: 0.22; nsp3: 0.77; nsp5: 0.70; nsp7: 0.77: nsp12: 0.88; nsp14: 0.68; nsp15: 0.76). These proteins are predicted to be involved in evading host immune defenses, in enhancing viral expression and in cleavage of the replicase polyprotein, based on prior studies of related betacoronaviruses^11,12^. Hence, this lower mutation rate may indicate purification selection to maintain these functions essential for efficient immune evasion and subsequent viral dissemination. Indeed, structural proteins in coronaviruses undergo a greater degree of antigenic variation which increases the fitness of the virus by means of adaptation to the host and by facilitating immune escape^13^.

Structural protein modeling confirmed that most of the nonsynonymous mutations in the nonstructural proteins were neutral (Supplementary Figure 5-13). Conversely, several nonsynonymous mutations in the spike protein might have functional consequences: notably, the G clade–defining mutation D614G is located in subdomain 1 (SD1; Figure 5, Supplementary Figure 5). In the trimeric S, D612 engages stabilising interactions within SD1 (R646 or the backbone of F592, depending on the chain) and with the S1 of the adjacent chain (T859 and K854). Replacement of D614 with a glycine would entail losing these stabilising electrostatic interactions and increase the dynamics in this region. Notably, V483A (found in 13 cases in Washington state, USA), V367F (in 6 cases: 5 in Paris, France; 1 in Hong Kong) and G476S (in 7 cases: 4 in Washington state, USA; 1 in Idaho state, USA; 1 in Oregon state, USA; and 1 in Braine-l’Alleud, Belgium) are localised in the receptor binding domain (RBD) of the spike protein which mediates binding to the host receptor angiotensin-converting enzyme 2 (ACE2) (Figure 5)^3^. All of the viral genomes harbouring V483A and G476S mutations belong to Clade S. Interestingly, the V367F mutation has appeared independently in Clade V and Clade S, suggesting that this mutation contributes to viral fitness. We found the V483A substitution in 13 closely-related cases from Washington State. An equivalent amino acid substitution located in a similar position within the RBD in the MERS-CoV spike protein reduces its binding to its cognate receptor DPP4/CD26^14^ (Supplementary Table 2). However, V483 is more than 10Å away from ACE2 and could affect receptor binding by SARS-CoV-2 only indirectly by altering the structural dynamics of the RBD loop it is a part of. The V367F mutation is located in an even greater distance from ACE2 (Figure 5, Supplementary Figure 5D). The exchange of the small hydrophobic residue valine with a bulky hydrophobic phenylalanine might influence the efficiency of glycosylation of the nearby N343, or the positioning of the sugars. The substitution G476S would lead to possible clashes with predicted interacting ACE2 residues and with the RBD residue N487. However, minor reorientation of the side chains might allow an additional hydrogen bond to be formed between S476 and ACE2 Q24 and E23, thus enhancing the affinity (Supplementary Figure 5D) to ACE2. An equivalent amino acid substitution located in an analogous position within the RBD in the SARS-CoV spike protein was associated with neutralisation escape from monoclonal antibodies, together with other mutations observed previously^15^ (Supplementary Table 2). Given that V483A and V367F are solvent exposed and markedly alter the surface characteristics of the RBD, they might also facilitate antibody evasion. Escape mutations in the RBD of the SARS-CoV S protein (T332I, F460C and L443R) were identified previously^15^. These mutations negatively impact viral fitness through reducing the affinity to the host receptor^16^ (Supplementary Table 2). The nonsynonymous mutations in the N protein, which have key roles in viral assembly, might also have functional implications. The hotspot mutations S202N, R203K and G204R all cluster in a linker region where they might potentially enhance RNA binding and alter the response to serine phosphorylation events (Supplementary Figure 6). The clade I–defining mutation in the nucleocapsid protein, which has key roles in viral assembly, is synonymous. However, we observed nonsynonymous mutations in the nucleocapsid protein that are predicted to have functional implications. The hotspot mutations S202N, R203K and G204R all cluster in a linker region where they might potentially enhance RNA binding and alter the response to serine phosphorylation events (Supplementary Figure 6).

**Figure 5:**
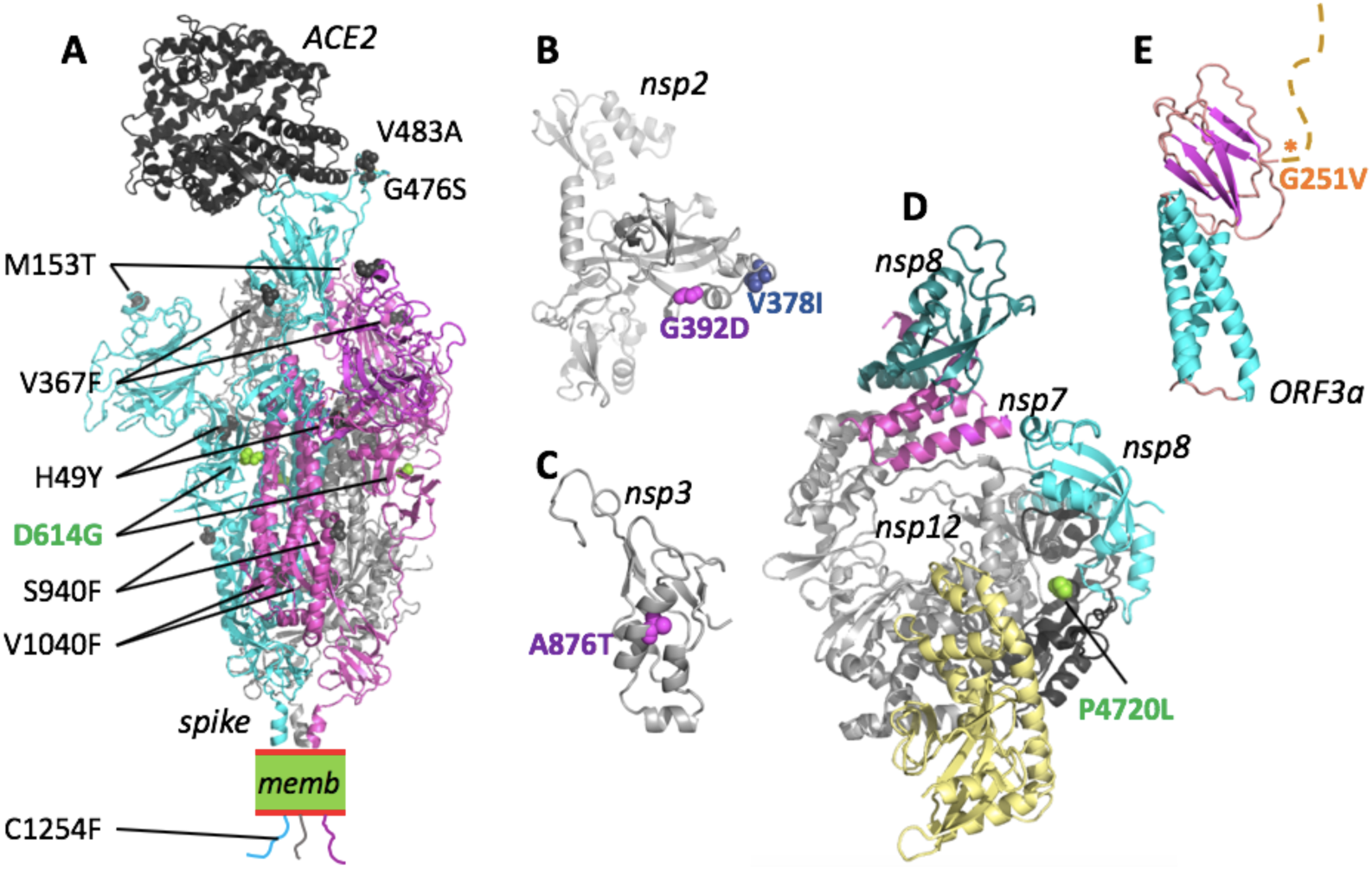
Mapping of SARS-CoV-2 clade-defining mutations onto the proteins. Nonsynonymous mutations for proteins where the 3D structure was experimentally determined (spike, nsp12/7/8) or can be inferred with reasonable confidence. Mutations are colour-coloured as for the corresponding clades in Figure 3 (D: magenta; G: light green; I: blue; V: orange). For a detailed analysis, see Supplementary Figures 5-13. (A) The structure of the SARS-CoV-2 spike trimer in its open conformation (chains are cyan, magenta and grey) bound to the human receptor ACE2 (black) modeled based on PDB accessions 6m17 and 6vyb. Identified nonsynonymous mutations are shown as spheres in the model. For reasons of visibility only mutations of two of the three spike chains are labeled. memb. indicates the plasma membrane. (B) Fragment comprising residues 180-534 of nsp2, modelled by AlphaFold^35^. Both clade-defining mutations are located in solvent-exposed regions and would not lead to steric clashes. (C) The substitution A876T (corresponding to residue A58 in the nsp3 cleavage product numbering) is situated in the N-terminal ubiquitin-like domain of nsp3. The structure of this domain can be inferred based on the 79% identical structure of residues 1-112 from SARS-CoV (PDB id 2idy). The substitution A876T can be accommodated with only minor structural adjustments and is not expected to have a substantial influence on the proteins stability or function. (D) The structure shows the nsp12 in complex with nsp7 (magenta) and nsp8 (cyan and teal), based on PDB 7btf. P4720 (P323 in nsp12 numbering) is located in the ‘interface domain’ (black). In this position, the P323L substitution is not predicted to disrupt the folding or protein interactions and hence is not expected to have strong effects. (E) A theoretical model for the Orf3a monomer has been proposed by AlphaFold^36^. The structure-function relationship of this protein remains to be clarified. The mutation G251V is located C-terminal to the β-sandwich domain and the tail (marked by an asterisk).

## DISCUSSION

In this study, we have defined 5 major clades (G, I, S, D and V) and 9 minor clades (H2, L2, G1110A, H, Y, C17373A, S2, I2 and K) which covers ∼89% of the 2,058 high quality genomes available until March 31^st^ in GISAID database. The clustering of these genomes revealed the spread of clades to diverse geographical regions (Figure 1, Figure 3). This pattern contrasts with those observed for other epidemic coronaviruses, such as MERS-CoV, which display distinct geographical clustering^17^. For example, clade G, which was first detected on January 28^th^, has reached 40 countries and 130 cities, within a span of 10 weeks (Supplementary Table 1). This pattern demonstrates efficient viral transmission through frequent intercontinental travel during the period when international travel restrictions were only present sporadically, which has enabled the virus to spread to multiple distant locations within a short period of time. This observation reinforces the importance of curtailing international travel and imposing restrictions early in pandemics and imposing social distancing in order to contain the global spread of viruses.

We have observed a distinct distribution of the major clades in different parts of the world (Figure 3). Most of the viral genomes that have not been assigned to a major clade are found in Asia and have earlier detection times in January and February at the start of the epidemic in China (Supplementary Figure 2). We observed a decrease in the genetic diversity of the virus over time following dissemination from China, especially in Europe and North America that each notably now has a predominant clade type, which we believe to be associated with a founder effect whereby a single clade was introduced and subsequently disseminated (Figure 3). An important caveat of the present study is that the current sampling of available public genomes does likely not represent the extant genetic diversity of virus populations in circulation due to biases of genome data deposits from the sequencing laboratories based mainly in the northern hemisphere and new datasets may define new clades in the near future from regions, including Africa, the Indian subcontinent and Latin America with comparably few genomes available at present. In this case, additional identifiers within an evolving barcode scheme can be added to track and monitor future emerging clades with higher resolution. On the other hand, the genetic stability of SARS-CoV-2^18^ may result in the continuing circulation of a limited number of clades until such time as mitigation measures including the isolation of vulnerable populations and the availability of efficacious antivirals and vaccines reduces the genetic diversity in circulation. This molecular genotyping approach has been demonstrated for other viruses (e.g. measles, poliovirus, rotavirus and human papillomaviruses) with herd-immunity vaccination programmes working to eliminate pathogens from endemic circulation in humans^19,20,21,22^. The availability of a barcoding scheme that rapidly generates a SNP allowing clade assignment will be critical in this elimination phase when widespread availability of vaccines permits eradication of SARS-CoV-2 from endemicity in humans.

Our work provides a baseline global genomic epidemiology of SARS-CoV-2 prior to introduction of therapeutic and prophylactic approaches. The mutational landscape of global populations of over SARS-CoV-2 6,000 genomes provides an evidence-based framework for tracking the clades that comprise the pandemic on different continents. However, due to the bias in the representation of countries depositing the SARS-CoV-2 genomes with over-representation of North American and European genomes (28.3% and 47.2% respectively) and the available genome data representing only a minute proportion of the total COVID-19 positive cases from each of these regions (America, 0.27%; Europe, 0.31%; China, 0.31%. Data collected from GISAID^9^ and COVID-19 dashboard^1^), the genetic barcode described hererin may need to be updated in order to be globally representative, once sufficient numbers of genomes covering less represented parts of the world are eventually sequenced and deposited to publicly-available database. We envisage a qPCR-based allelic discrimination approach, such as PCR allele competitive extension (PACE), which would enable rapid turnaround in real-time following the identification of a laboratory-confirmed case. This would allow viral genetic epidemiological data to be added to contact tracing information to allow efficient detection of circulating SARS-CoV-2 clades that will allow discrimination of autochthonous and imported clades to aid progressive elimination of the genetic diversity and, ultimately, eradication in all regions. A robust genetic barcoding scheme for SARS-CoV-2 can facilitate this molecular tracking of larger numbers of laboratory-confirmed cases and by implementing such a facile genotyping approach upstream of next-generation sequencing will allow whole genome sequencing to be performed on selected cases. This is of particular relevance when the available genomes represent only a small sample of the over 2.5 million total COVID-19 cases globally to date.

## METHODS

### Phylogenomic Analysis

1,427 coronavirus genomes were downloaded from Virus Pathogen Database and Analysis Resource (ViPR)^23^ on February 14^th^ 2020, including 329 SARS, 35 SARS-CoV-2, 61 NL63, 521 MERS, 52 HKU1, 170 OC43, 97 bovine coronaviruses and 61 mouse hepatitis viruses. Sequence alignment was performed using MAFFT(version 7.407)^24^ software and then trimmed by trimAL(version 1.4.1)^25^. Phylogenetic analyses of the complete genomes were done with FastTree(version 2.1.10)^26^ software with default parameters, and iTOL(version 5)^27^ was used for phylogenetic tree visualisation.

2,127 complete SARS-CoV2 genomes were downloaded from the GISAID9 (March 31th 2020). 69 of those genomes were removed due to poor assembly quality resulting in 2,058 complete genomes that were subsequently used for analysis. Sequence alignment was performed with MAFFT(version 7.407)^24^ and trimmed by trimAL(version 1.4.1)^25^. Phylogenetic analyses of the complete genomes were performed with RAxML^28^ (version 8.2.12) with 1,000 bootstrap replicates, employing the general time-reversible nucleotide substitution model. iTOL(version 5)^27^ was used for the phylogenetic tree visualisation. SNPs from each of the genomes were called by Parsnp (version 1.2) from the Harvest suite^29^ using MN908947.3 as the reference genome, and the SNPs were further annotated by SnpEff(version 4.3m)^30^. The monoclades associated with SNPs with a frequency ≥40 were defined as major clades and monoclades associated with SNPs with a frequency ≥5 were defined as minor clades.

### Protein Structural Analysis

Experimentally determined protein structures were obtained from the Protein Data Bank (PDB). SwissModel^31^, I-Tasser^32^, RaptorX^33^ and an in-house modelling pipeline was used to produce protein structure homology models. Phobius^34^ was used for prediction of trans-membrane regions. RaptorX was also used for predicting secondary structure, protein disorder and solvent exposure of amino acids. Pymol(version 1.8.6.2) was used for visualization.

## ACKNOWLEDGEMENTS

This work was supported by funding from King Abdullah University of Science and Technology (KAUST), Office of Sponsored Research (OSR), under award number FCC/1/1976-25-01. Work in AP’s laboratory is supported by the KAUST faculty baseline fund (BAS/1/1020-01-01) and research grants from the Office for Sponsored Research (OSR-2015-CRG4-2610, OCRF-2014-CRG3-2267). We thank all laboratories which have contributed sequences to the GISAID database. We thank Olga Douvropoulou, Raeece Naeem Mohamed Ghazzali and Sharif Hala for their support during the work. We also thank Richard Culleton (Nagasaki University, Japan) and Gabo Gonzalez (UCD, Ireland) for their critical comments on the manuscript draft.

## AUTHOR CONTRIBUTIONS

AP conceived the study and supervised the work; AP and QG designed the analysis. QG, MS and SA performed the data analysis and prepared the initial draft of the manuscript, followed by edits from AP, MC and RN. All authors have commented on various sections of the manuscript.

## SUPPLEMENTARY FIGURES

**Supplementary Figure 1.**
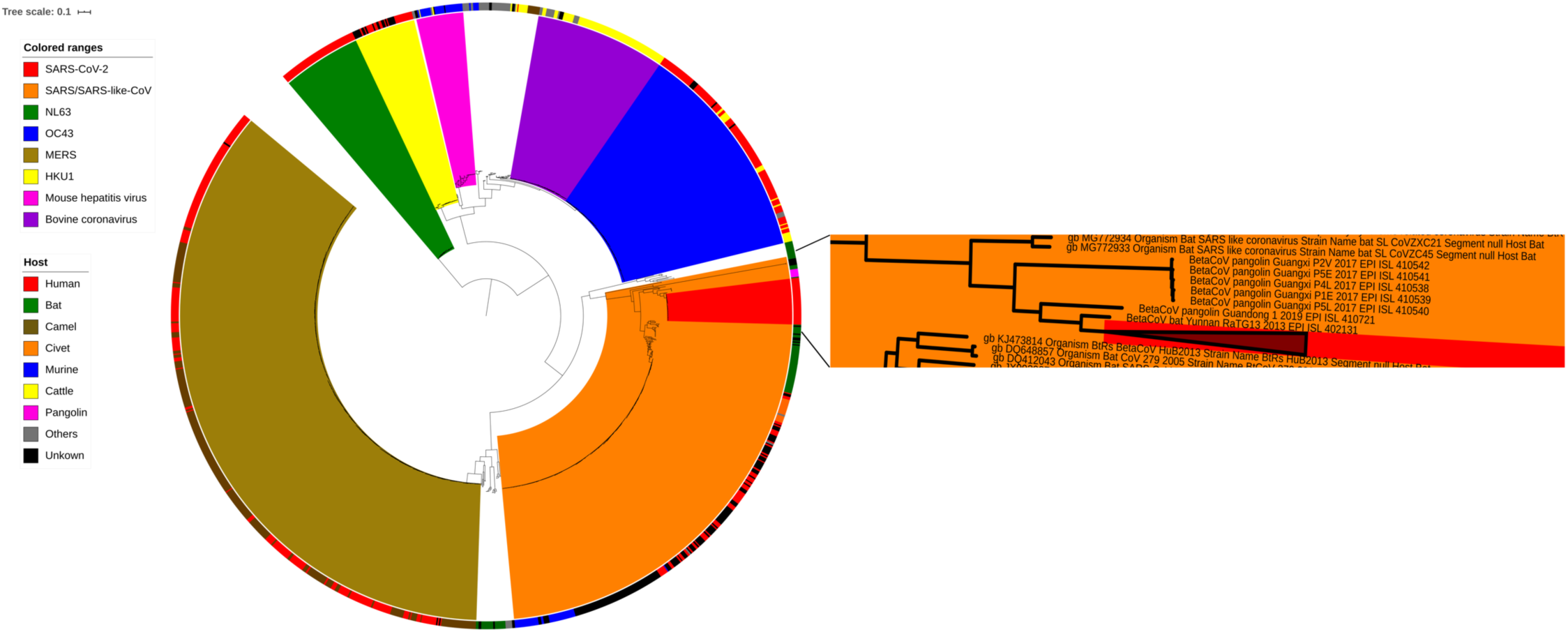
A maximum likelihood phylogenomic tree based on whole genomes from SARS-CoV-2 and other coronaviruses. All available complete genomes of an alphacoronavirus (NL63) and several major groups of betacoronavirus species were used to construct the phylogenetic tree. The colour-coded outer circle represents the host of each specific genome sequence. The SARS-CoV-2 part of the phylogenetic tree has been expanded for better resolution of the tree with closely related viral species. A total of 1,427 SARS-CoV-2 whole genomes were illustrated in the phylogeny.

**Supplementary Figure 2:**
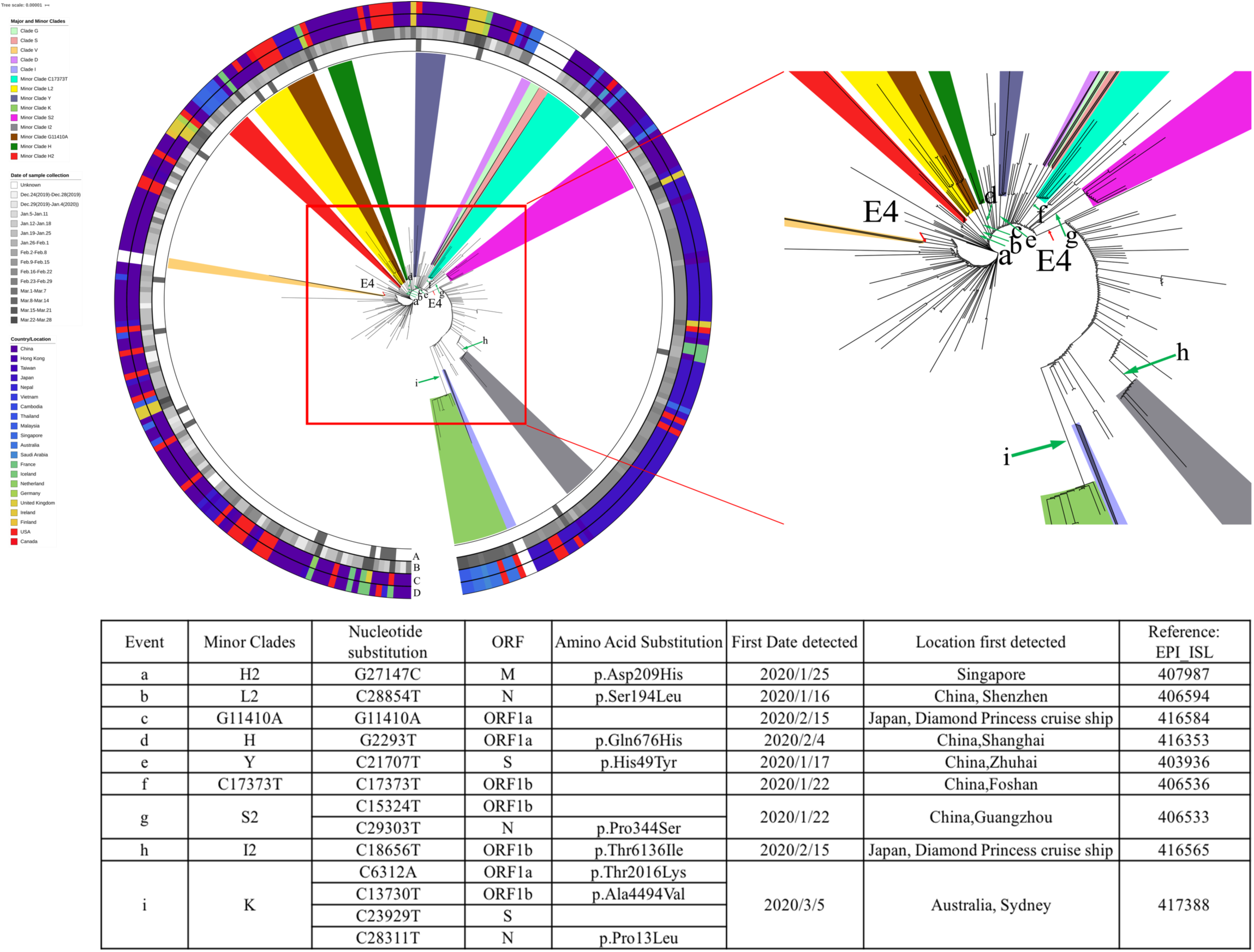
Minor clades SARS-CoV-2. For illustration purposes, the major clades have been collapsed and the central part of the figure has been enlarged. Most of the genomes that cannot be assigned to any major clade are derived from Asian countries indicative of high genetic diversity in the relatively early stages of the local epidemics and that a founder effect likely explains the observations of single predominant clades in North America and Europe which exhibit a reduction in genetic diversity. These genomes obtained were further analysed and assigned to minor clades. A minor clade was defined with n≥5 cases of SARS-CoV-2 based on phylogenetic and SNP analysis and the events associated with them are indicated by the green arrow. The information contained in the concentric circles: A, the imported cases; B, the collection date of each genome at one-week resolution; C, collection locations of the genomes; D exposure locations of the genomes.

**Supplementary Figure 3.**
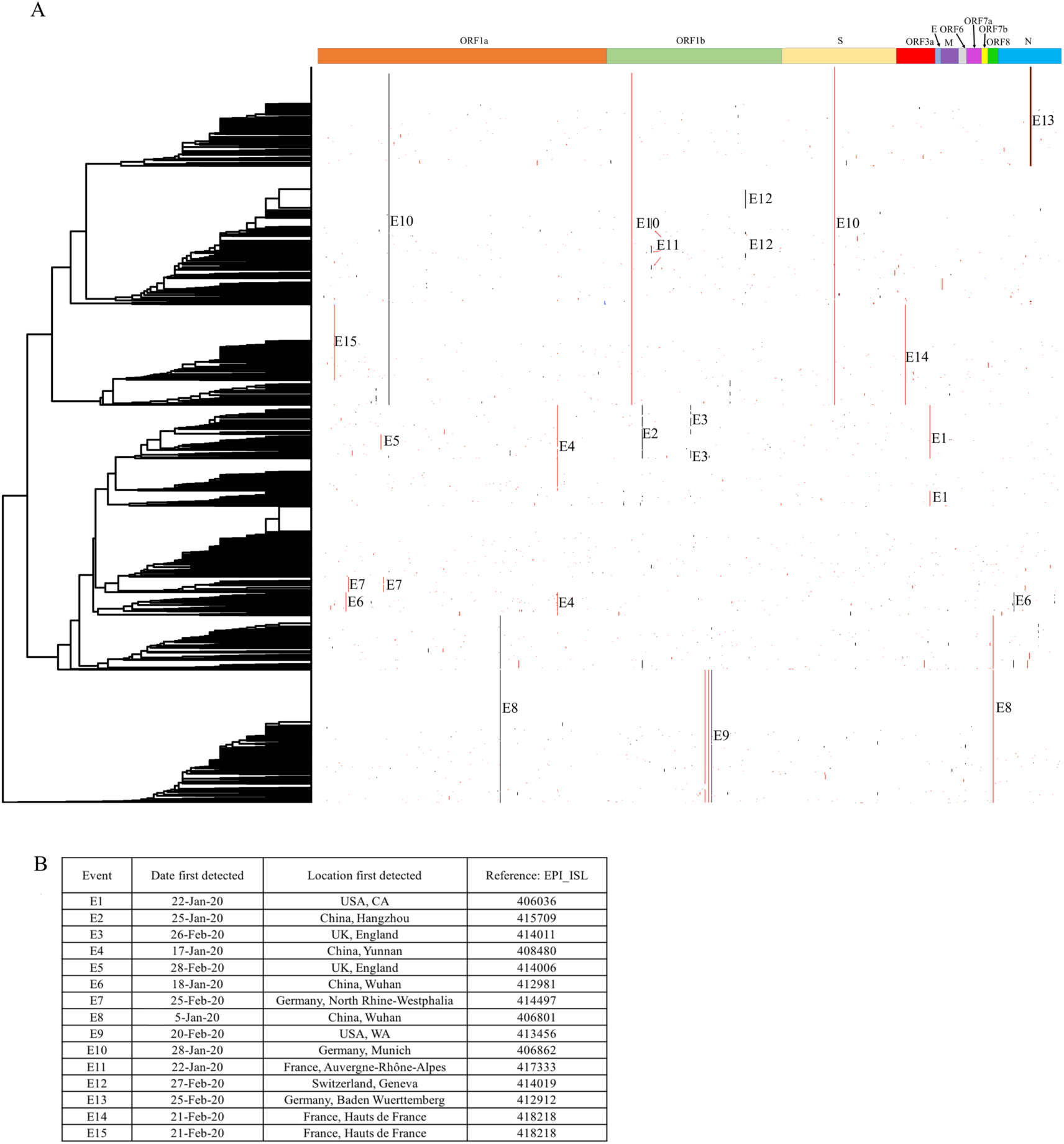
The genomic position and details of SNPs in 2,058 SARS-CoV-2 genomes and the amino acid substitution eventss. (A) Nonsynonymous mutations are marked in red, and synonymous mutations are labeled in black. (B) The first appearance and the location of each event

**Supplementary Figure 4.**
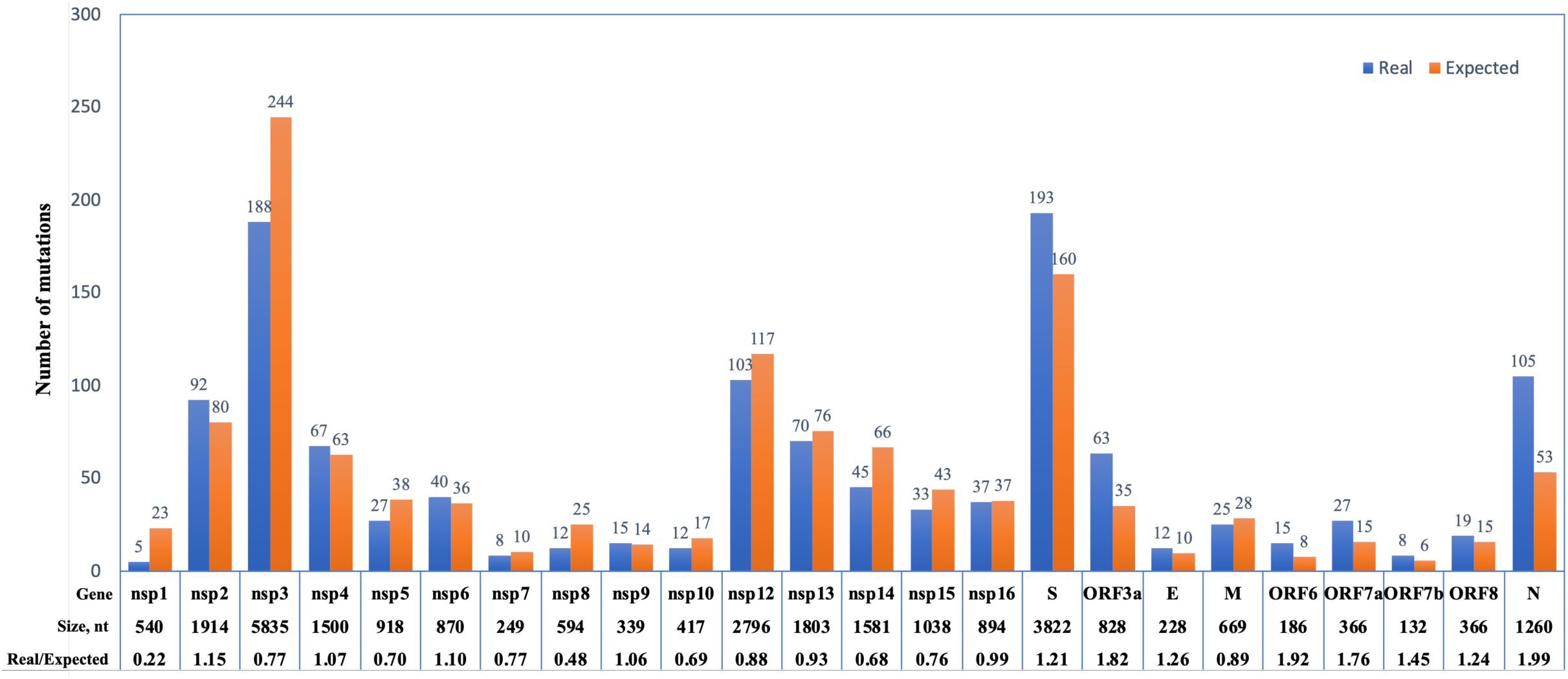
Number of unique mutations in the genome of SARS-CoV-2. The number of mutations (blue) were taken from 2,058 available genomes from GISAID (March 31^st^ 2020). The number of expected mutations (orange) were calculated based on the assumption that there are no purifying or selection pressures present against different categories of point mutations, and all of the mutations occurring randomly according to the size of each gene.

**Supplementary Figure 5.**
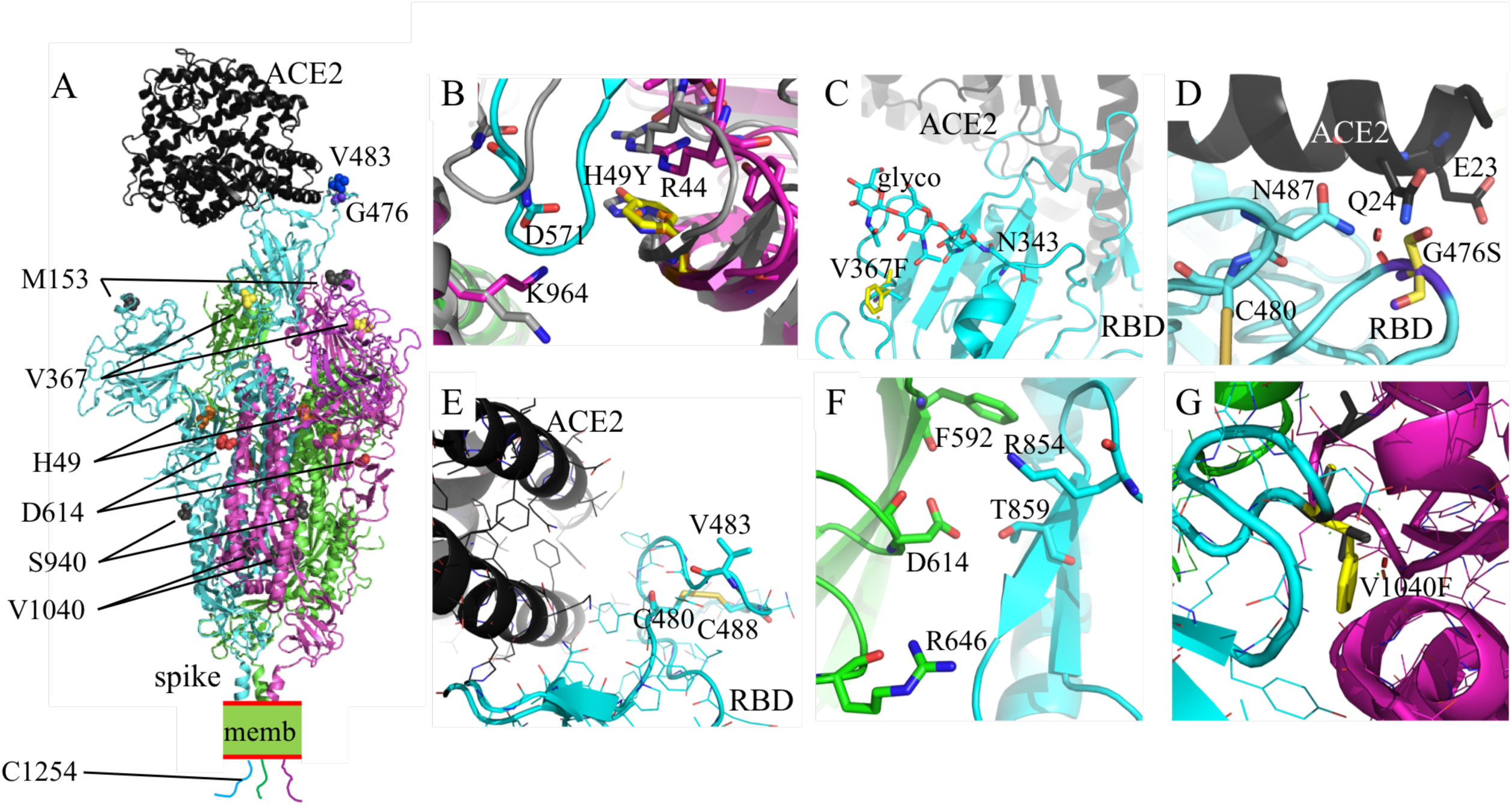
Mutations in the SARS-CoV-2 spike protein. (A) An overview of an S trimer. The structure of the SARS-CoV-2 S trimer in its open conformation (chains are cyan, magenta and green) bound to the human receptor ACE2 (black) has been modeled based on PDB accessions 6m17 and 6vyb. Identified nonsynonymous mutations are shown as sphere models. For reasons of visibility only mutations of two of the three S chains are labeled. Memb indicates the membrane. The cytoplasmic tails are depicted in the same colours as the three spike chains. B-E) Magnified regions of the mutations. (B) H49Y: H49 is located in the N-terminal domain (NTD) which is in contact with the sub-domain 2 (SD2) of a symmetry-related molecule. The relative position of NTD and SD2 changes considerably upon going from the closed (grey coloured) to the open state (cyan and magenta). H49 stabilises the rim of the NTD through cation-π-stacking with R44. A tyrosine in position 49 would be able to perform the same role and does not lead to clashes. However, the substitution may slightly alter the stability of this interaction and the interaction between the NTD and SD2, which, in turn, might have subtle effects on the stability and equilibrium of open and closed conformations of S. (C) V367F: V367 is part of the receptor binding domain (RBD), however located too far away from the ACE2-binding site to directly affect receptor binding. V367 is surface exposed, and its substitution would not create clashes with other protein regions. However, the exchange of a small with a bulky hydrophobic residue would alter the surface characteristics of this region, which might influence the efficiency of glycosylation (stick model) of the nearby N343, or the positioning of the sugars. Additionally, the altered RBD surface could potentially interfere with antibody recognition. (D) G476S: G476 is located in the RBD. It is positioned solvent-exposed in a SARS-CoV-2-specific loop. This loop is stabilised by a disulphate bridge (C480:C488; C480 is shown as stick model). In the open, ACE2-bound conformation of the RBD, G476 is close to ACE2 Q24 and E23. The substitution G476S would lead to light clashes with these ACE2 residues (indicated as red spheres) and with the RBD residues N487. However, minor reorientation of the side chains might allow an additional hydrogen bond to be formed between S476 and ACE2 Q24 and E23, thus enhancing the affinity. (E) V483A: V483 is also located in the RBD, solvent-exposed in the same SARS-CoV-2-specific loop as G476 (C480:C488 are shown as stick models). In the open, ACE2-bound conformation of the RBD, V483 is more than 10Å away from the ACE2 receptor, and hence does not contribute to direct binding or stability. In the closed conformation, this loop is not modeled in the EM structures (PDB 6vyb), inferring it is flexible in the absence of ACE2. Superimposition of the ACE2-bound conformation of the RBD onto RBDs in a closed conformation shows that this loop region would stick out into the solvent. Substitution of V483 is consequently not predicted to have a strong impact on receptor binding or protein stability. By lowering the hydrophobic surface, this substitution might however reduce the non-specific stickiness of this loop region, and/or affect binding of antibodies. (F) D614G: D614 is located in the SD1. In the trimeric S, D612 engages stabilising interactions within the SD1 (R646 or the backbone of F592, depending on the chain) and with the S1 of the adjacent chain (T859 and K854). Replacement of D614 with a glycine would entail losing these stabilising interactions and increase the dynamics in this region. (G) V1040F: V1040 is located in a loop region that makes hydrophobic contacts between stalk regions of the spike trimer. The V1040F substitution is possible without steric hindrance, and would slightly increase the hydrophobic contacts between the chains. The other mutations are not shown in detail, but are evaluated as follows. M153T: M153 is located in the NTD in a solvent-exposed loop. N electron density was modeled for this loop in the cryo-EM structure (6vyb) suggesting it is flexible. The substitution is expected to be neutral. S940F: S940 is located in the stalk region, in a solvent-exposed turn. Introducing the bulkier phenylalanine in this position would not destabilise the structure but locally change the surface characteristics. C1254F: C1254 is the last cysteine in a cysteine-rich unstructured short cytoplasmic region. This region is required for efficient membrane fusion. The exact mechanism remains to be elucidated, and hence we cannot assess the exact impact of the C1254F mutation.

**Supplementary Figure 6.**
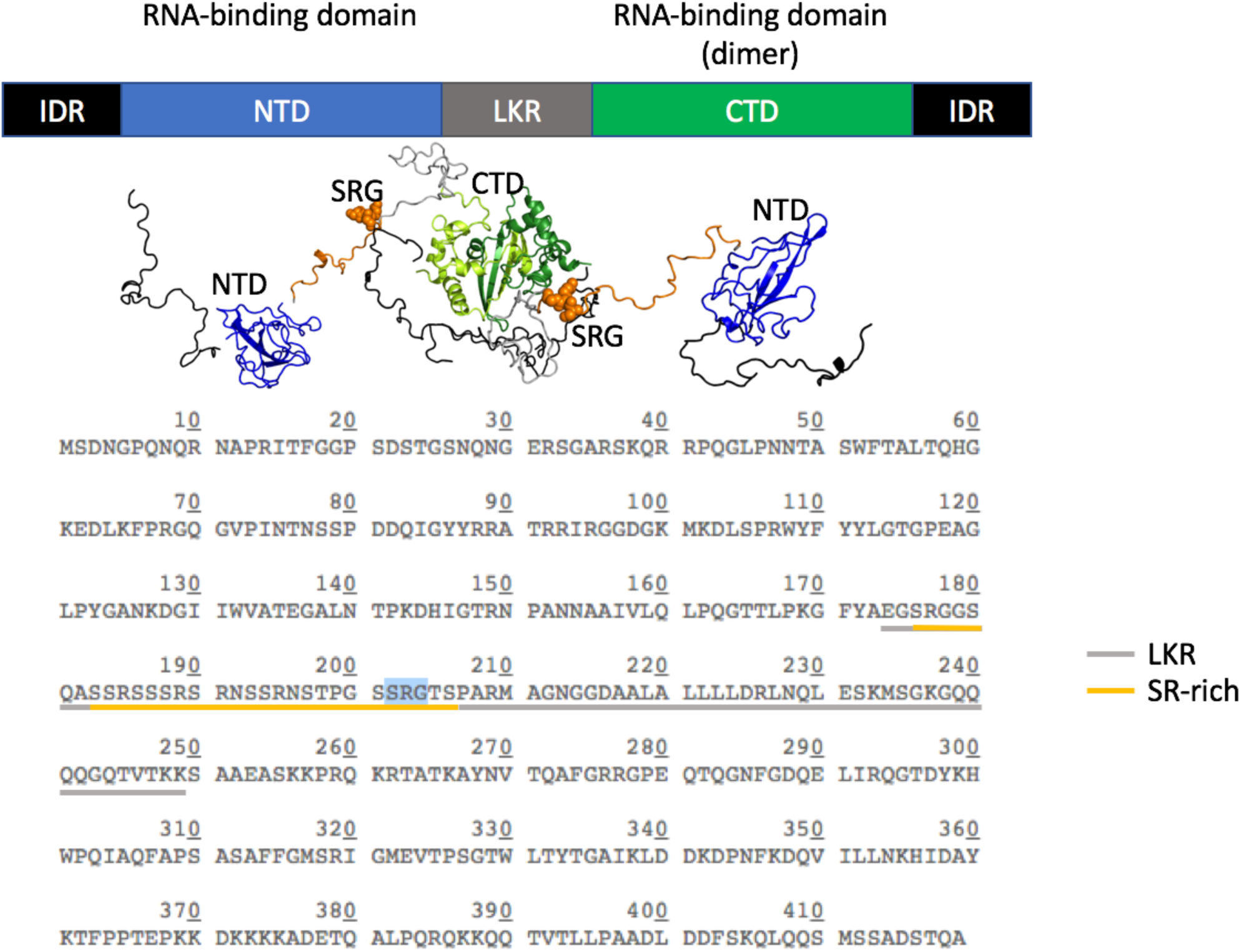
Mutations in the SARS-CoV-2 nucleocapsid. The SARS-CoV-2 nucleoprotein (N) compacts the viral genome into a helical ribonucleocapsid and hence has key roles in viral assembly^1^. N contains two folded domains, the N-terminal and the dimeric C-terminal domain (NTD; blue, and CTD; green, respectively). Both domains are flanked by flexible regions, namely a flexible linker between them (LKR, underlined in the bottom panel sequence) and ∼40 residue intrinsically disordered regions (IDRs) as tails. The NTD and CTD are RNA-binding regions, but the LKR and IDRs also affect RNA-binding of N^2^. The LKR contains a serine-arginine (SR)-rich region, which contributes to RNA binding and N oligomerization. The arginines may promote RNA-binding through electrostatic interactions, whereas the serines are putative phosphorylation sites that would counteract RNA binding and possibly favour oligomerisation upon phosphorylation^3^. The hotspot mutations S202N, R203K and G204R are all within the SR region of the LKR. S202N would delete a putative phosphorylation site. Given the large number of alternative serine phosphorylation sites, and the presence of many asparagines within the linker, this S/N substitution might not have a very strong influence. The homologous substitution R203K is also expected to have only minor effects. G204R might increase binding to RNA, and/or affect other homo- or heterologous interactions. In summary, the substitutions might potentially enhance RNA binding and alter the response to serine phosphorylation events. The structural figure consists of homology models for the NTD and CTD, linked by arbitrary but stereochemically-plausible linker regions.

**Supplementary Figure 7.**
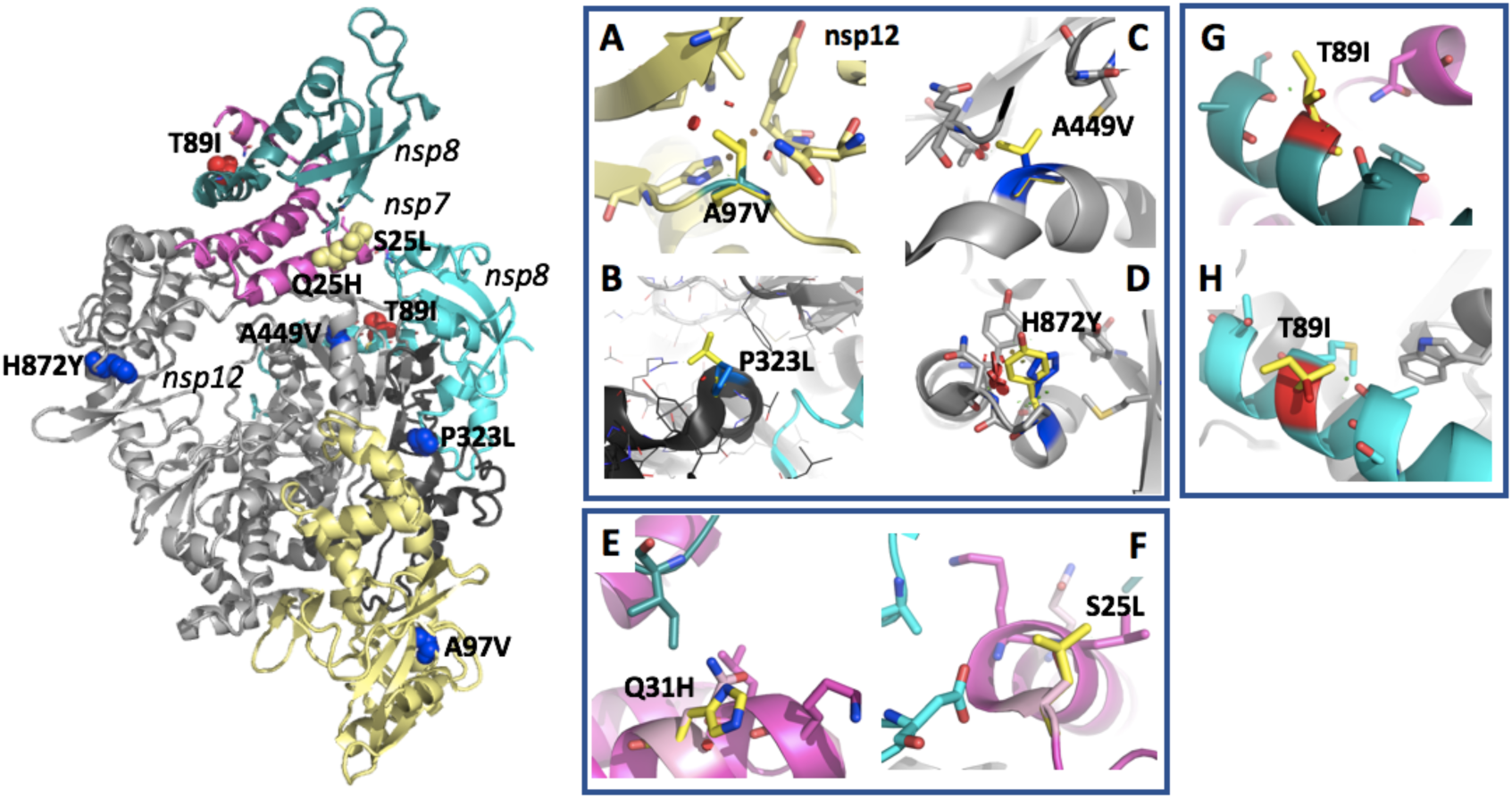
Mutations in SARS-CoV-2 *ORF1ab*, nsp12: The nsp12 RNA-dependent RNA polymerase forms a complex with one nsp7 and two nsp8, which markedly enhances its polymerase activity^4^. The shown SARS-CoV-2 nsp12 structure is composed of a nidovirus-unique N-terminal extension (pale yellow), a linker domain (black) and the RNA-dependent RNA polymerase (RdRp) domain (grey). The structure shows the nsp12 in complex with nsp7 (magenta) and nsp8 (cyan and teal), based on PDB 7btf. (A) A4494 (A97 in nsp12 numbering; shown as a blue sphere models) is located in the N-terminal extension of the polymerase^5,6^. A4494 is sealing the hydrophobic core of the N-terminal lobe, and its side chain is not solvent exposed. Its replacement with the hydrophobic but slightly bigger valine (yellow) only leads to minor clashes (small red discs) that would not have a significant impact on the function or stability. (B) P4720 (P323 in nsp12 numbering) is located in the ‘interface domain’ (black). In this position, the P323L substitution (yellow) is not predicted to disrupt the folding or protein interactions and hence is not expected to have strong effects. (C) A4846V (residue A449 in nsp12 numbering; blue) is located in the finger domain, with its side chain pointing inwards, contributing to a hydrophobic interaction with the adjacent beta strand. The substitution of A4846 by a valine (yellow) is tolerated in this context. Leading only to minor clashes, it might slightly improve the stability of this region. (D) H5269 (H872 in nsp12 numbering; blue) is located in the thumb domain. It is at the tip of a solvent exposed turn, where its replacement by a tyrosine (yellow) would be tolerated without functional impact. **Nsp7**: S2884 and Q3890 (S25 and Q31 in nsp7 numbering; both in light yellow) are solvent exposed on a helix that makes contact with both nsp8 and nsp12. (E) S25 is capping the N-terminal end of this helix. Its substitution with a leucine (yellow) does not cause steric problems, but would lead to loss of the capping hydrogen bond. However, D163 from one of the nsp8 molecules also performs a capping function in the complex, and hence S25L would only have a slightly destabilising effect. (F) Q31 is located on the surface of the helix. Although Q31 is close to nsp8, its substitution with a histidine (yellow) would not influence this interaction measurably. **Nsp8**: Only T4031 (T89I in nsp8 numbering; red) is included in the structural model. In both nsp8 molecules, T89 is located solvent exposed on a helix. In neither of the two nsp8 molecules would the substitution T89I create steric problem or affect the interaction with nsp12 or nsp7.

**Supplementary Figure 8.**
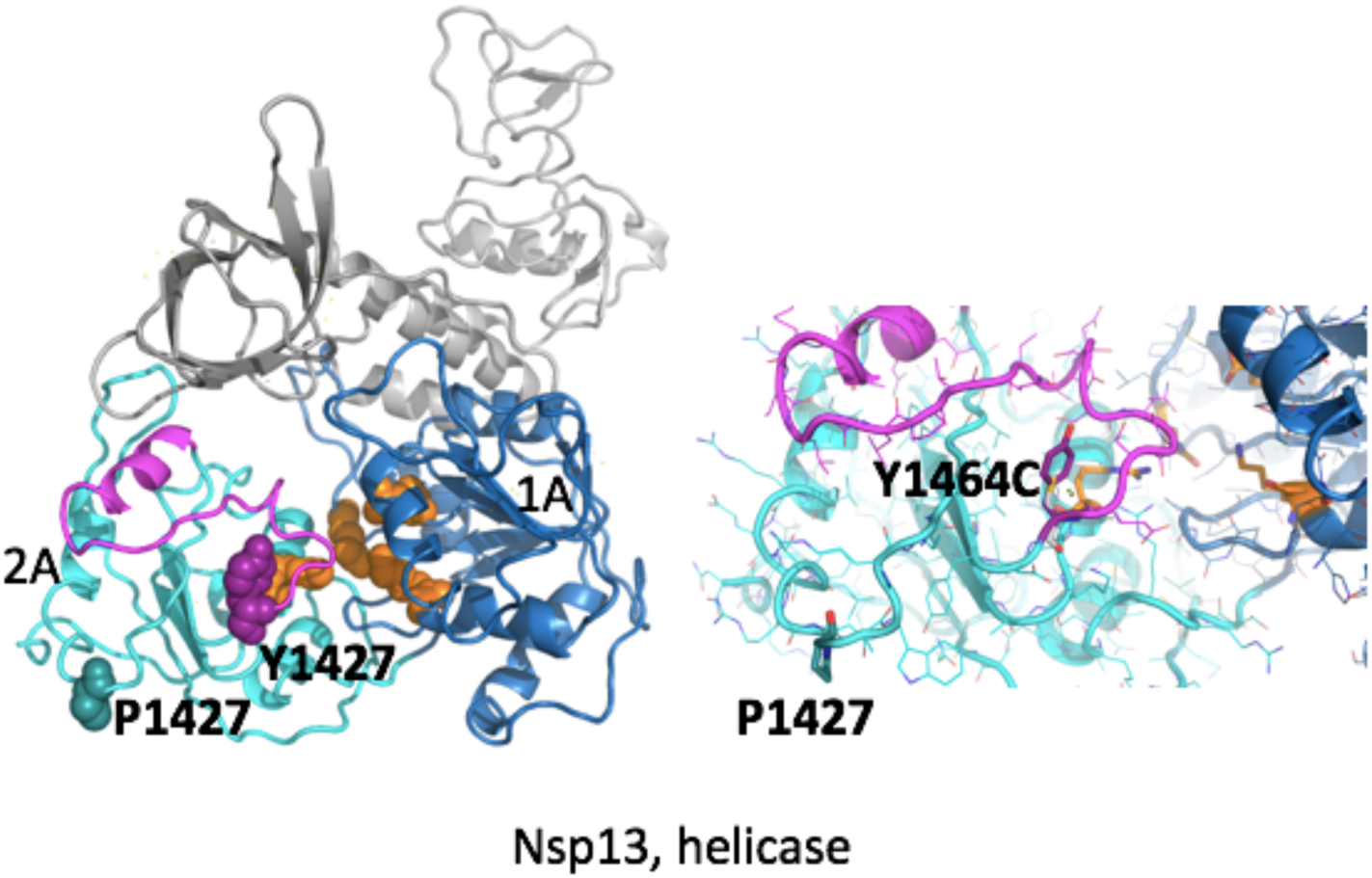
Mutations in SARS-CoV-2 *ORF1ab*, nsp13: P1427L and Y1464C are both in the helicase nsp13, which catalyses the unwinding of duplex oligonucleotides into single strands. Both residues are located on the same side of the 2A domain of the helicase. The 2A and adjacent 1A domains coordinate together to complete the final unwinding process. The helicase has been modelled based on the 99.8% identical SARS nsp13 (PDB id 6jyt). The domains 1A and 2A are coloured in blue and cyan, respectively. The residues involved in NTP binding are shown in orange. Residues on domain 2A that are involved in RNA binding are shown in magenta. The other domains are grey. *Left*: overview of the complete structure. Key residues are shown as sphere models. *Right*: zoom into the mutated area. Key residues are shown as stick models. For Y1464 the *in silico* mutated cysteine is shown in white. P1427 is located in a solvent-exposed loop region that has not yet reported to be involved in nucleotide binding. Its substitution with a leucine is not expected to create noticeable effects. Y1464 is part of a region that contributes to binding and unwinding of duplex oligonucleotides^7^. Its substitution by a cysteine would decrease the stability and enhance the dynamics of this particular region, and might affect RNA binding and processing.

**Supplementary Figure 9.**
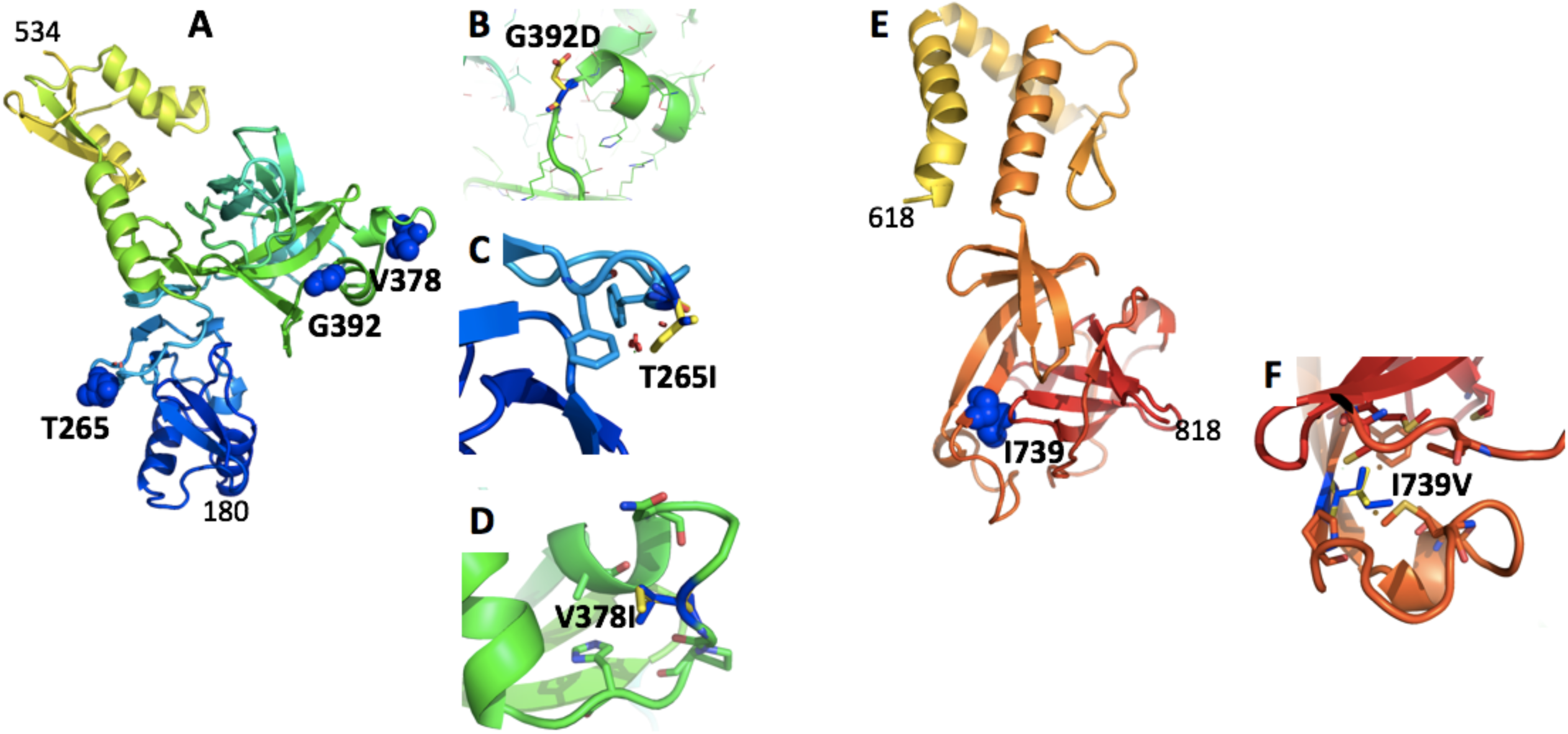
Mutations in SARS-CoV-2 *ORF1a*: nsp2. No experimental structure of sufficiently close homologue is known for nsp2. This structural analysis is therefore based on the proposed theoretical AlphaFold model. (A) Shown is a model for residues 180-534, colour ramped from blue to yellow. T265 and G392 are shown as blue sphere models. (B) T265 (blue stick model, corresponding to residue T58 in the nsp2 cleavage product numbering) is located at the tip of a loop that is pinned to the core of the N-terminal domain (blue) via hydrophobic residues (two phenylalanines are shown as stick models). Being exposed to the solvent, the T265I substitution (shown in yellow) is not expected to have significant impact on protein fold or function. (C) G392 (G212 in nsp2 numbering; shown in blue) is placed in a solvent accessible loop, according to the AlphaFold model. In this position even the non-conservative substitution with an aspartic acid (yellow) is not predicted to have significant impact on protein fold or function. (D) V378 is located in a surface exposed loop. Its substitution by an isoleucine will not create steric clashes, nor lead to loss of hydrophobic interactions. (E) A second nsp2 fragment is shown, colour ramped from yellow to red, comprising residues 618 to 818. I793 is shown as blue sphere model. (F) I739V (I559 in nsp2 numbering; shown in blue) is part of a hydrophobic core of a small C-terminal domain. Its substitution with an only slightly smaller hydrophobic valine (yellow) would only have insignificant destabilising effects on this region.

**Supplementary Figure 10.**
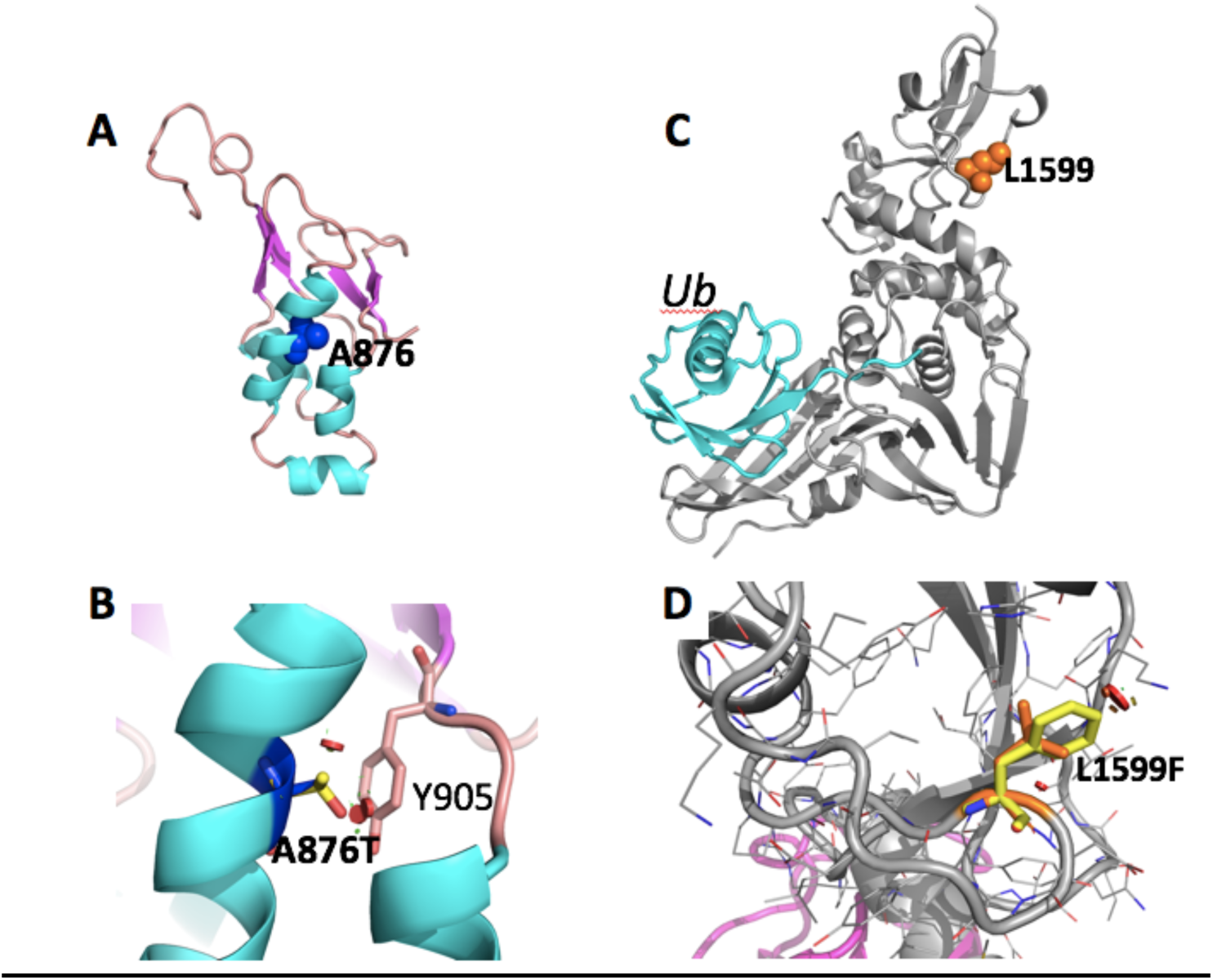
Mutations in SARS-CoV-2 *ORF1a*: nsp3. The substitution A876T (corresponding to residue A58 in the nsp3 cleavage product numbering) is situated in the N-terminal ubiquitin-like domain of nsp3. (A) The structure of this domain can be inferred based on the 79% identical structure of residues 1-112 from SARS-CoV (PDB id 2idy). Helices are coloured in cyan, strands in magenta and loops in pale orange. A876 is shown as blue sphere model. (B) Zoom onto A876. A876 (blue stick model) is placed within a helix, engaging hydrophobic contacts with Y905 (stick model). The substitution into a slightly larger threonine (yellow stick model) can be accommodated with only minor structural adjustments (clashes are shown as red spheres) and hence the A876T mutation is not expected to have a substantial influence on the proteins stability and function. (C) L1599F is located in the papain-like protease. The structure (grey) has been modelled based on the ∼83% identical SARS-CoV structures in complex with ubiquitin (Ub, cyan, based on 4m0w). L1599 is remote from the active site in the Ubl domain. (D) Its substitution does not lead to clashes and is expected to be neutral in terms of function and protein stability. No templates have been identified for V378I.

**Supplementary Figure 11.**
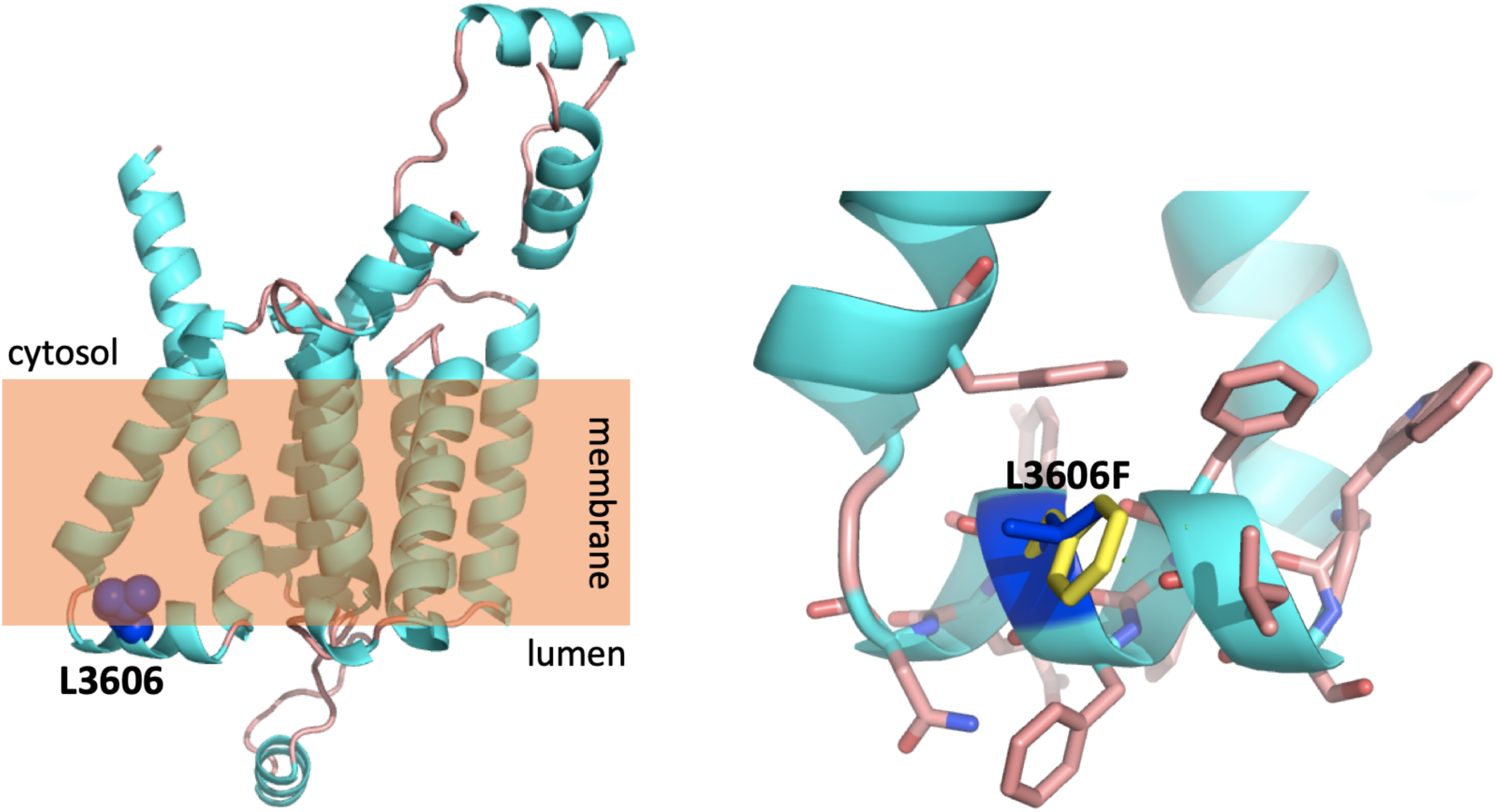
Mutations in SARS-CoV-2 *ORF1a*: nsp6. L3606 (corresponding to residue L37 in the nsp6 cleavage product) is a multi-pass transmembrane protein for which no experimental structure of sufficiently close homologue is known. This structural analysis is therefore based on the proposed theoretical AlphaFold model (*Left*). Helices are colored in cyan, loops in pale orange. L3606 is shown as blue sphere model. The endoplasmic reticulum membrane is shown in orange; lumen and cytoplasmic sides are indicated. *Right*: close-up view of the location of L3606 (shown as blue stick model). L3606 is located in a predicted helical region that is partly submerged in the membrane, lying parallel to its luminal surface. According to the structural model, L3606 is exposed to the membrane, and hence its substitution with a larger but still hydrophobic phenylalanine would not impact structure or function.

**Supplementary Figure 12.**
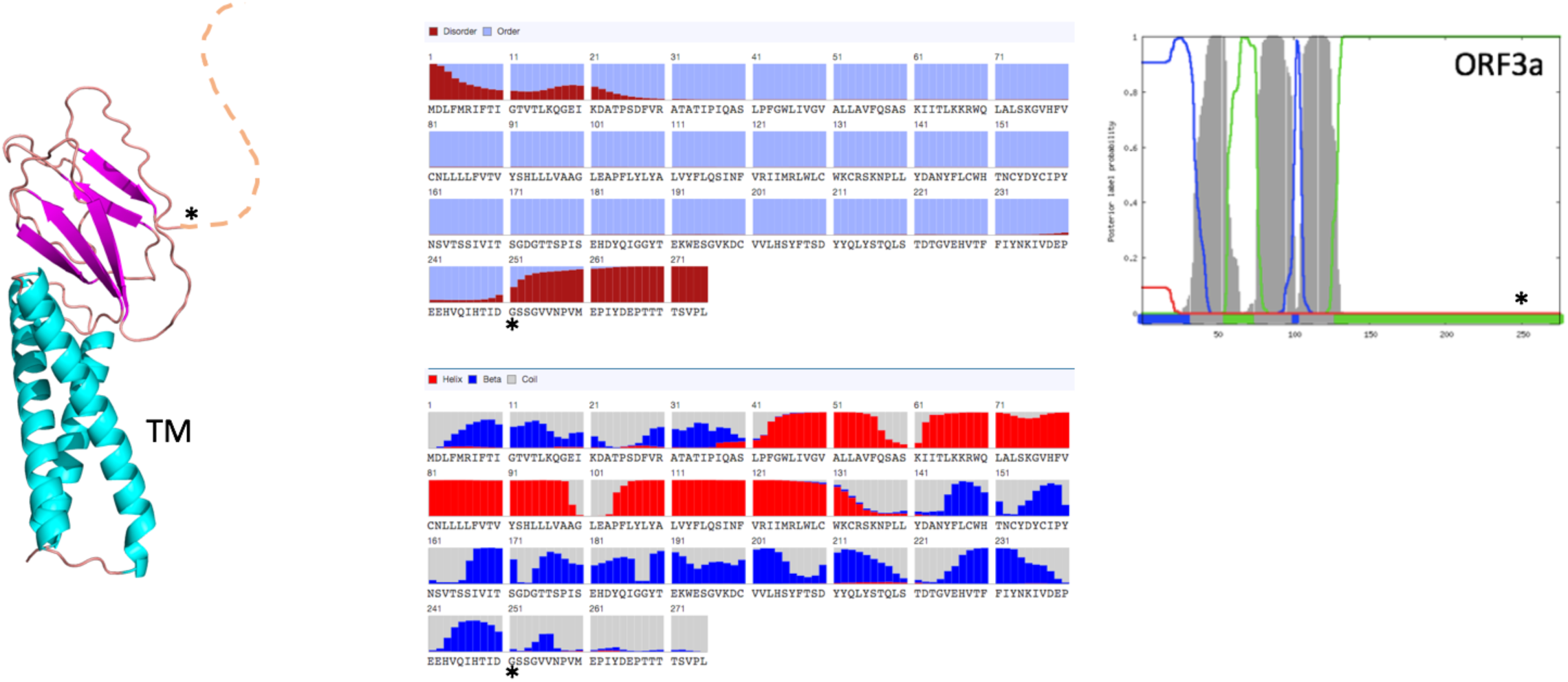
Mutations in SARS-CoV-2 *ORF3a*. ORF3a is a viroporin that forms a pentameric potassium-sensitive ion channel. ORF3a activates the inflammasome which facilitates viral release and aggravates disease symptoms^8^. ORF3a contains a 3-pass α-helical TM region (cyan in left panel, and greyed regions in TM prediction, right panel) and a domain predicted to have a β-sandwich fold (magenta; see also middle panel and right panel). The ORF3a N- and C-terminal tails are predicted to be disordered (middle panel, top). A theoretical model for the Orf3a monomer has been proposed by AlphaFold^9^. The structure-function relationship of this protein remains to be clarified. The mutation G251V is located C-terminal to the β-sandwich domain and the tail (marked by an asterisk). In this position, the substitution is not expected to affect the protein fold or function significantly.

**Supplementary Figure 13.**
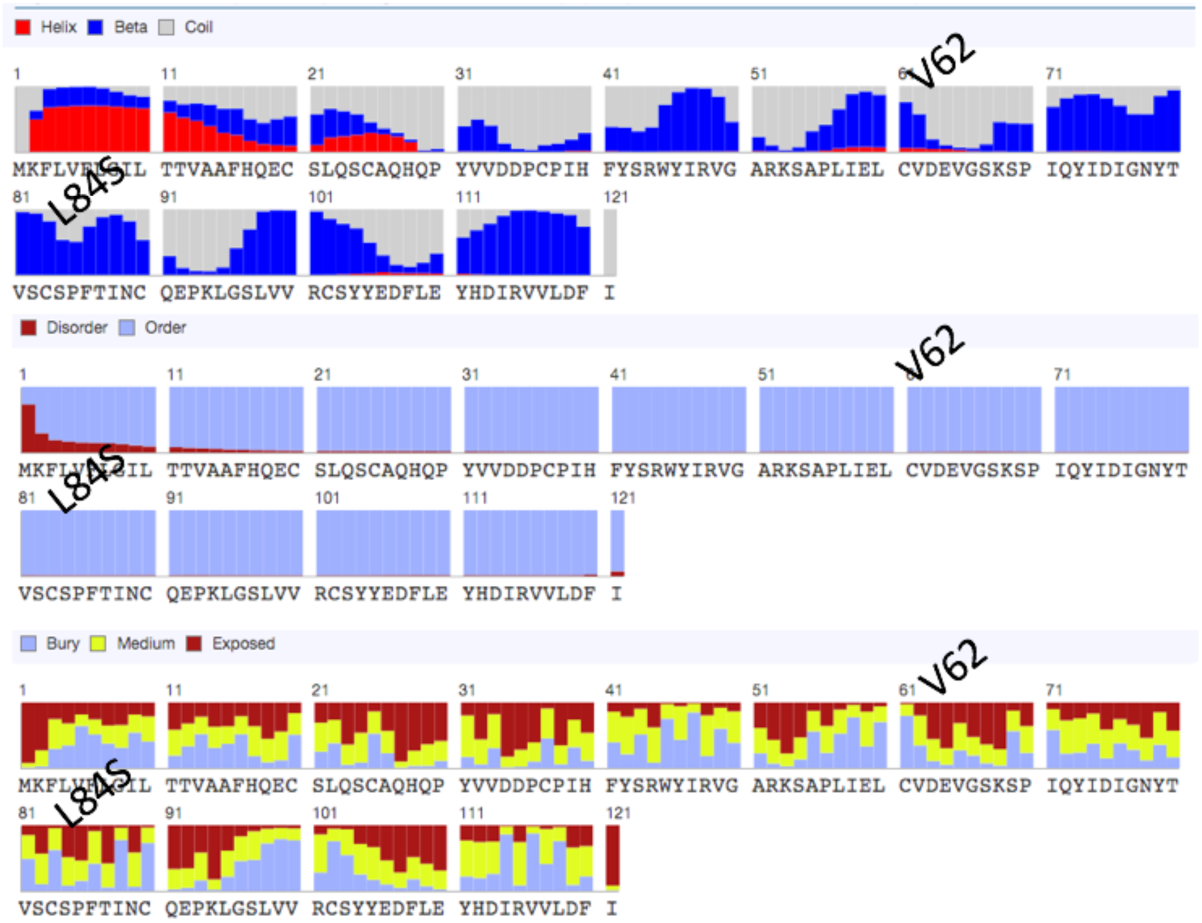
Mutations in SARS-CoV-2 *ORF8*. ORF8 has an N-terminal *sec*-pathway signal peptide with a cleavage site after residue 15, suggesting that it is secreted into the extracellular space. Following signal peptide cleavage, the ORF8 protein core is predicted to consist largely of β-strands and features seven cysteines (top panel). We predict that this protein adopts a cysteine disulfide-bond stabilised β-sandwich structure inferring that ORF8 also functions as a ligand binding module. Homology modeling or *ab initio* servers failed to produce a model consistent with the proteins’ secondary structure predictions. V62 and L84 are predicted to be partially solvent-exposed (bottom), and located at the end (V62) or in the middle (L84) of secondary structural elements (depending on the prediction server, the region surrounding V62 is predicted as strand or helix; L84 is in a β strand). Hence, we can only speculate that the mutations V62L and L84S might have little effect on the stability of the disulphate-stabilised fold, but might have minor (or no) effects on ligand binding.

## SUPPLEMENTARY TABLES

**Supplementary Table 2.** Experimentally validated amino acid substitutions resulting in escape mutants or having functional significance in N and S proteins of SARS-CoV and MERS-CoV.

| Protein | Residue | Virus | Substitution in<br>SARS-CoV-2 | GISAID (EPI_ISL) | Functional role of the residue | Reference |
| --- | --- | --- | --- | --- | --- | --- |
| N | T149 | SARS-CoV | T148I | 406595 | T149 is an epitope that interacts with neutralising antibody. | 10 |
|  | K250 | SARS-CoV | K249I | 408515 | K250 is an essential motif for binding SN5-25 antibody. | 10 |
|  | S328 | SARS-CoV | S327L | 413557 | S328 forms inter-dimer disulfide bonds connecting the two<br>β-strands. | 11 |
| S | R395 | SARS-CoV | R408I | 413522 | R395 interacts with one of the most potent neutralising<br>antibody m396. | 12 |
|  | D463 | SARS-CoV | G476S | 417085, 417380, 418055,<br>418077, 417353, 41708, 416447 | D463G is associated with neutralisation escape from MAbs<br>S224.1. | 13 |
|  | I529 | MERS-CoV | V483A | 414618, 415596, 415605,<br>417072, 417073, 417075,<br>417076, 417111, 417139,<br>417159, 417160, 418029, 418071 | I529T reduces affinity of RBD to the DPP4/CD26 receptor. | 14 |

